# Assessing the coupling between local neural activity and global connectivity fluctuations: Application to human intracranial EEG during a cognitive task

**DOI:** 10.1101/2021.06.25.449912

**Authors:** Manel Vila-Vidal, Mariam Khawaja, Mar Carreño, Pedro Roldán, Jordi Rumià, Antonio Donaire, Gustavo Deco, Adrià Tauste Campo

**Author notes:** These authors jointly supervised this work (Gustavo Deco, Adrià Tauste Campo). Corresponding author. Address: Center for Brain and Cognition, Department of Information and Communication Technologies, Universitat Pompeu Fabra, 08005, Barcelona, Spain.

## Abstract

Cognitive-relevant information is processed by different brain areas that cooperate to eventually produce a response. The relationship between local activity and global brain states during such processes, however, remains for the most part unexplored. To address this question, we designed a simple face-recognition task performed in patients with drug-resistant epilepsy and monitored with intracranial EEG. Based on our observations, we developed a novel analytical framework (named “local-global” framework) to statistically correlate the brain activity in every recorded gray-matter region with the widespread connectivity fluctuations as proxy to identify concurrent local activations and global brain phenomena that may plausibly reflect a common functional network during cognition. The application of the local-global framework to the data from 3 subjects showed that similar connectivity fluctuations found across patients were mainly coupled to the local activity of brain areas involved in face information processing. In particular, our findings provide preliminary evidence that the reported global measures might be a novel signature of functional brain activity reorganization when a stimulus is processed in a task context regardless of the specific recorded areas.

**Data availability statement:** Due to institutional restrictions, the data that supports the findings of this study can be accessed only with a data sharing agreement. All code used in this work can be found at https://github.com/mvilavidal/localglobal2022.

**Funding statement:** MVV was supported by a fellowship from ”la Caixa” Foundation, Spain (ID 100010434, fellowship code LCF/BQ/DE17/11600022). MVV and ATC were supported by the Bial Foundation grant 106/18. GD and ATC were supported by the project ”Clúster Emergent del Cervell Humà” (CECH, ref. 001-P-001682), within the framework of the European Research Development Fund Operational Program of Catalonia 2014-2020. GD was supported by a Spanish national research project (ref. PID2019-105772GB-I00 MCIU AEI) funded by the Spanish Ministry of Science, Innovation and Universities (MCIU), State Research Agency (AEI); HBP SGA3 Human Brain Project Specific Grant Agreement 3 (grant agreement no. 945539), funded by the EU H2020 FET Flagship programme; SGR Research Support Group support (ref. 2017 SGR 1545), funded by the Catalan Agency for Management of University and Research Grants (AGAUR); Neurotwin Digital twins for model-driven non-invasive electrical brain stimulation (grant agreement ID: 101017716) funded by the EU H2020 FET Proactive programme; euSNN European School of Network Neuroscience (grant agreement ID: 860563) funded by the EU H2020 MSCA-ITN Innovative Training Networks; Brain-Connects: Brain Connectivity during Stroke Recovery and Rehabilitation (id. 201725.33) funded by the Fundacio La Marato TV3; Corticity, FLAG–ERA JTC 2017, (ref. PCI2018-092891) funded by the Spanish Ministry of Science, Innovation and Universities (MCIU), State Research Agency (AEI).

**Conflict of interest disclosure:** The authors declare no conflicts of interest.

**Ethics approval statement:** The study was conducted in accordance with the Declaration of Helsinki. All diagnostic, surgical and experimental procedures have been previously approved by The Clinical Ethical Committee of Hospital Clínic (Barcelona, Spain). In particular, the specific proposal to run the cognitive experiments for this study was approved in March 2020 under the code number HCB/2020/0182.

**Patient consent statement:** Informed consent was explicitly obtained from all participants prior to the recordings and the performance of the tasks.

## 1 Introduction

Human cognition implies the contribution of different brain areas that interact to process incoming information and eventually produce a response. A classical localist approach in cognitive neuroscience has attempted to assign cognitive functions to specific brain areas and understand their role at different stages of the cognitive process. This approach has proven successful in partially explaining a number of cognitive processes such as attention, memory or decision making with different recording modalities (Hubel and Wiesel, 1959; Brunel and Wang, 2001; Wang, 2002; Deco and Rolls, 2005; Kahana, 2006; Lachaux et al., 2012). In sharp contrast, more recent studies have taken a globalist approach, focusing on brain states that can be measured by means of statistical dependencies across the whole brain (Sporns et al., 2005; Axmacher et al., 2008; Bressler and Menon, 2010; Palva et al., 2010; Bassett et al., 2011; Wang et al., 2015; Deco et al., 2015; Cruzat et al., 2018). Yet, a fundamental question linking the two perspectives remains for the most part unaddressed: how does the processing of cognitive-relevant information in each functionally involved brain area relate to the brain’s global state?

To tackle this question, we designed a simple face-recognition paradigm that patients with drug-resistant epilepsy conducted during the pre-surgical intracranial monitoring period (Munari and Bancaud, 1985; Kahane et al., 2003; Lachaux et al., 2003; Engel et al., 2005). During this procedure, the intracranial activity of up to 200 electrode contacts in varied regions from cortical and subcortical regions is simultaneously recorded, which sheds light into the mechanisms of neural activity associated with consciously perceiving and reporting a visual stimulus. In particular, we collected intracranial EEG data (also often referred in the literature as local field potentials, LFP) from depth electrodes stereotactically implanted for pre-surgical diagnosis in 3 drug-resistant epilepsy patients while they were performing the cognitive task.

Inspired by the recorded data, we developed and systematized a methodological pipeline integrating neuroanatomic information, clinical reports, signal processing functions and statistical analysis, with the aim to localize and quantify time-varying human neural activity in the context of a cognitive task. High-frequency LFP power is known to display high correlation with spiking activity of local neuronal assemblies (Hipp et al., 2012; Pesaran et al., 2018). Based on these results, we used LFP high-frequency power activations as proxy for locally generated activity. Yet, the specific frequency range of such activity can vary depending on the type of LFP recording technique. To avoid making further assumptions, we adopted a data-driven approach and defined the range depending on the observed activations. In order to characterize global network states, we resorted to functional connectivity analysis. Activity in the beta band and below is known to display long-range coherence and it is thought to have a more widespread origin (Pesaran et al., 2018), reflecting possible concurrent inputs or more global states. Based on these premises, we defined two independent functions in the low-frequency range to measure the brain sites’ connectivity consistency across time and across trials. In addition, we took advantage of the referential montage to capture these global cofluctuations, as suggested by previous literature (Tauste Campo et al., 2018).As a result of our work, we propose a novel data-analysis framework (named “local-global” framework) that statistically correlates the brain activity in every gray-matter recording site with the predefined connectivity functions to assess the coupling between local neural activations and the brain’s global connectivity during cognition. We first applied this framework to the data gathered from 2 patients with epilepsy conducting the same task paradigm, showing that global connectivity fluctuations were temporally associated with variations of local activity in task-relevant areas. Finally, we validated our methodology in a third subject with a slightly different task paradigm confirming most of our previous findings.

## 2 Methods

### 2.1 Ethics statement

The study was conducted in accordance with the Declaration of Helsinki and informed consent was explicitly obtained from all participants prior to the recordings and the performance of the tasks. All diagnostic, surgical and experimental procedures have been previously approved by The Clinical Ethical Committee of Hospital Clínic (Barcelona, Spain). In particular, the specific proposal to run the cognitive experiments for this study was approved in March 2020 under the code number HCB/2020/0182.

### 2.2 Participants and behavioural task

Intraranial EEG recordings during performance of certain tasks were acquired in 2 subjects with pharmacoresistant epilepsy during the diagnostic monitoring period in Hospital Clínic (Barcelona, Spain). Details on patients’ demographic information is given in Table 1. The two participants had normal or corrected-to-normal vision. The task was designed to characterize brain responses to static face recognition. Stimulus presentation was designed and responses were collected using Psychtoolbox 3 for Matlab.

**Table 1:** Demographic data. F: female; M: male; FCD: focal cortical dysplasia

| Subject | Sex | Age (years) | Epilepsy onset (years) | Suspected epileptic focus | Hand laterality | Implantation | Number of electrodes (right/left) | Number of contacts (right/left) | Number of contacts included in analysis |
| --- | --- | --- | --- | --- | --- | --- | --- | --- | --- |
| 1 | F | 51 | 27 | Right lateral frontal / insula (FCD) | Right | Right | 16/0 | 175/0 | 47 |
| 2 | M | 49 | 15 | Left dorso-medial frontal | Right | Left | 0/7 | 0/93 | 26 |

Subjects viewed *N* (64 and 71 for subjects 1 and 2, respectively) face images of different identities on a laptop screen. Approximately one half of these images were familiar faces and the remaining half were images from people that the subject was unlikely to know. Familiar faces were selected after a short interview with the subject in which they introduced themselves and talked about their hobbies and interests. Potentially unfamiliar faces were extracted from an extensive database and were selected according to the age, country and background of the subject. Face images had a resolution of 160×160 pixels, and were presented at the centre of the screen framed by thin lines that randomly changed their color (green and red) from image to image presentation to help the subject maintain attention. Each trial (Fig. 1A) started with a pre-stimulus blank screen that lasted ITI = 0.5 ms (inter-trial interval), then a face image was presented during time *T*_stim_ (1 and 0.9 s for subjects 1 and 2, respectively). Subjects were instructed to pay attention to the presented face image and then, after the face went off the screen, to respond during a maximum timeout of *T*_out_ (10 and 3 s for subjects 1 and 2, respectively) whether they recognized the specific person by pressing the trigger button of a joystick with the right hand. Either after pressing the joystick or when the maximum response timeout had elapsed, the next trial started as stated above. The parameters were adjusted depending on cognitive and experimental constraints associated with each subject. This task lasted in total between 5 and 12 minutes. Subject 1 experienced psychotic symptoms later on during the day of the task. We therefore decided to disregard all behavioral responses (recognized vs non-recognized) from our analysis in this subject.

**Figure 1:**
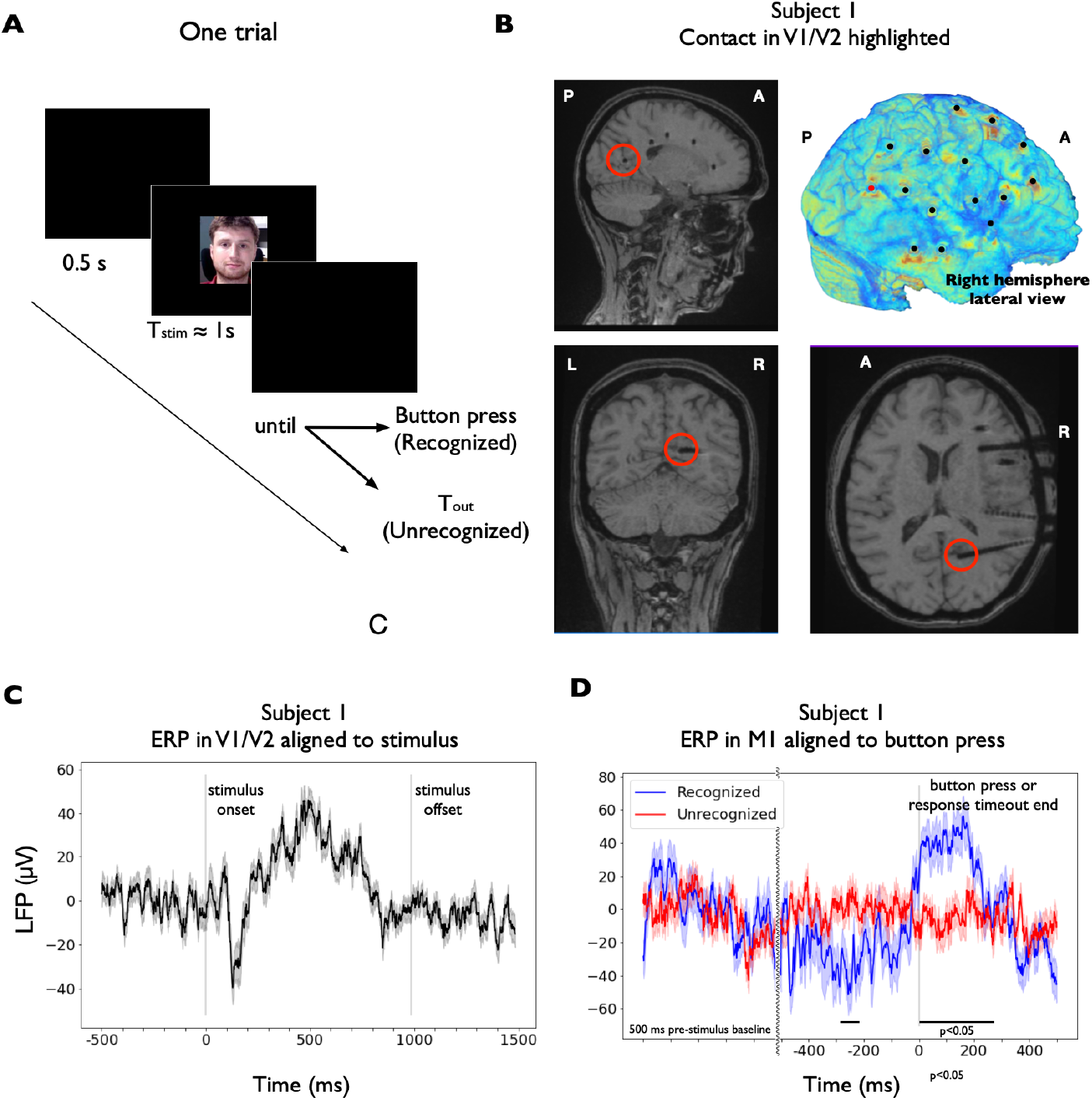
Experimental protocol, recording locations and neural responses. Experimental protocol, recording locations and neural responses. **(A)** Schematic representation of the behavioural task. Each trial consists of a pre-stimulus blank screen lasting 0.5 ms, a face image that remains present for approximately 1 s and a blank screen, where the subject has to report whether they recognized the person by pressing a trigger button. Alternatively, the trial ends after a maximum response timeout *T*_out_. **(B)** Subject 1 post-implantation MRI brain scans showing different electrode trajectories in sagittal, coronal and horizontal planes (from top left to bottom right) and 3D brain reconstruction (top right). The red circles highlight the trajectory of the electrode pointing towards the primary and secondary visual areas. **(C)** Event-related potential (ERP) in V1/V2 (deepest electrode contact of the trajectory highlighted in (B)) aligned to stimulus presentation (median ± SEM across N=64 trials). Signals were baseline-corrected on a trial-by-trial basis (baseline from 500 ms to 0 ms before stimulus presentation) before averaging. Vertical dark lines indicate the stimulus onset and offset times, respectively. **(D)** Event-related potential (ERP) in M1 (right-hand region) during recognized (button press with left hand, blue, median ± SEM across N=21 trials) and non-recognized (timeout end, red, median ± SEM across N=49 trials) trials aligned to button press or response timeout end, respectively (vertical dark line). Signals were baseline-corrected on a trial-by-trial basis (baseline from 500 ms to 0 ms before stimulus presentation) before averaging. Pre-stimulus baseline is also shown for comparison. Curvy line marks time discontinuity. In this case, we tested differences in the signals between both conditions. Black bars indicate time periods with significant differences between conditions (Ranksum test at each time point and across conditions, with a criterion of *P* < 0.05 for a minimum of 102 consecutive samples, 50 ms)

Across our initial study (subjects 1 and 2), we analyzed the data in two different settings: a stimulus-presentation-locked setting (subjects 1 and 2) and a motor-report-locked setting (subject 2). To further validate our methodology, we independently analyzed data from another subject (subject 3) that performed a cognitive task under a slightly different paradigm. Details about this patient can be found in Table S1. The task consisted of a total of 96 trials, with a similar structure to those of the main paradigm, which allowed for both stimulus and motor-locked analyses. In this case, however, each trial was preceded by a fixation cross that lasted 0.5 ms and that was meant to reset the subject’s attention. Although this difference might affect the cognitive processes involved in the task, the aim of this secondary analysis was to cross-validate some of our method’s assumptions and findings with a different dataset.

### 2.3 Data acquisition

LFPs were recorded using 16 and 7 (subjects 1 and 2, respectively) intracerebral multiple contact Microdeep® platinum-iridium Depth Electrodes (Dixi Medical, Besançon, France; diameter: 0.8 mm; 5 to 18 contacts, contact length: 2 mm, distance between contacts: 1.5 mm) that were stereotactically implanted using frameless stereotaxy, neuronavigation assisted, and intraoperative CT-guided O-Arm and the Vertek articulated passive arm. In total, 175, 93 and 146 contacts were implanted and recorded in subjects 1-3, respectively (see Tables 1 and S2 for details and Fig. 1B for an example of the implantation scheme of subject 1). The decision to implant, the selection of the electrode targets and the implantation duration were entirely made on clinical grounds using the standard procedure (Lachaux et al., 2003; Cardinale et al., 2013). All recordings were obtained using a standard clinical EEG system (XLTEK, subsidiary of Natus Medical) with a 2048 Hz sampling rate. All signals were referenced to the scalp electrode CPz.

Individual pre- and post-implant T1-weighted MR scans were used to determine contact localizations. MR scans were obtained with a 1.5 T unit (Magnetom Aera 1.5T; Siemens Medical Systems, Erlangen, Germany) with a specific protocol that included the following sequence: sagittal T1-weighted gradient recalled (repetition time [TR] 20 ms, echo time [TE] 7.38 ms, Flip Angle [FA] 20°, 1 mm slice thickness).

### 2.4 Anatomical localization of the SEEG electrode contacts and ROI definition

Contact anatomical locations were directly identified from the individual subjects’ post-implant MRI by visual inspection (MVV^1^), where contacts can be distinguished as dark spherical artifacts (diameter ≈ 3 mm; see Fig. 1B). MRI images were analyzed using the DICOM Viewer Osirix Lite (v.12.0.1). Contact localization within brain structures were therefore obtained with an error of the contact midpoint of approximately 1.5 mm. First, contacts were labelled either as grey matter (GM) or white matter (WM). Previous research suggests that electrical fields generated in GM can be measured by contacts in nearby WM up to ≈ 1 mm away (Buzsáki et al., 2012; Arnulfo et al., 2015; Narizzano et al., 2017). Based on this assumption, WM contacts lying at a distance below 1 mm from GM regions with no contact inside were assigned to that region and classified as GM. Contacts lying outside brain tissue or within altered brain tissue according to clinicians (e.g. heterotopias, focal cortical dysplasias) were excluded from the analysis. The electrode contacts lying in the suspected epileptic focus were identified by clinical experts using gold-standard procedures and were also excluded from the study.

GM contact locations were expressed in terms of the Desikan-Killiany (Desikan et al., 2006) brain atlas (34 ROIs per hemisphere), with an extra ROI for the hippocampus, by visually identifying well-defined anatomical landmarks (MVV). Table 2 summarizes the number of contacts and electrodes in each ROI for both subjects. See Table S2 for details about the third subject. Contacts were also approximately mapped to Brodmann areas, when possible. For further validation, automatic parcellations were created using the post-implant MRI scans and the Free-surfer software, which confirmed manual localizations. GM contacts were also mapped, when possible, to functional ROIs (regions of interest) as usually expressed in the cognitive literature based on fMRI and electrophysiological studies (e.g. V1/V2, DLPFC, VLPFC, M1, PMC), which were also confirmed by a neurologist (MK^2^).

**Table 2:** Implantation scheme. Regions of interest monitored in each patient are expressed in terms of the Desikan-Killiany atlas with an extra ROI for the hippocampus.

| ROI | P1 |  | P2 |  |
| --- | --- | --- | --- | --- |
|  | #Electrodes | #Contacts | #Electrodes | #Contacts |
| Rostral anterior cingulate | 1 | 1 | 2 | 3 |
| Caudal anterior cingulate | 1 | 2 | 1 | 1 |
| Posterior cingulate | 1 | 2 | 1 | 2 |
| Isthmus cingulate | 1 | 2 | 1 | 1 |
| Insula | 0 | 0 | 1 | 1 |
| Cuneus | 1 | 3 | 0 | 0 |
| Supramarginal | 2 | 7 | 0 | 0 |
| Inferior temporal | 1 | 1 | 0 | 0 |
| Middle temporal | 2 | 6 | 0 | 0 |
| Superior temporal | 2 | 8 | 2 | 6 |
| Superior frontal | 0 | 0 | 1 | 1 |
| Precentral | 1 | 3 | 1 | 3 |
| Pars triangularis | 0 | 0 | 1 | 3 |
| Caudal middle frontal | 1 | 3 | 0 | 0 |
| Rostral middle frontal | 2 | 5 | 2 | 5 |
| Hippocampus | 2 | 4 | 0 | 0 |
| **Total** | 18 | 47 | 13 | 26 |

For the purpose of this study, V1/V2 was defined roughly as the primary and/or secondary visual cortices. The DLPFC (dorsolateral prefrontal cortex) was defined roughly as the middle frontal gyrus, the VLPFC (ventrolateral prefrontal cortex) was defined as the inferior frontal gyrus and the superior parts of the pars triangularis, pars orbitalis and pars opercularis. M1 (primary motor cortex) was defined as the precentral gyrus and PMC (premotor cortex) was roughly defined as corresponding to Brodmann area 6, i.e., as a vertical strip extending from the cingulate sulcus to the lateral sulcus, including caudal portions of the superior frontal and middle frontal gyri, and rostrally bounded by the precentral gyrus. The remaining ROIs were referred to using its denomination in the Desikan-Killiany atlas. In particular, for this study: suparmarginal gyrus (SMG), insula (I), inferior temporal gyrus (ITG), middle temporal gyrus (MTG), superior temporal gyrus (STG), ventral anterior cingulate cortex (vACC), dorsal anterior cingulate cortex (dACC), and a single posterior cingulate cortex (PCC), under which we grouped the posterior cingulate and the isthmus cingulate.

### 2.5 Signal pre-processing

Besides the contacts mentioned in Section 2.4, we also excluded from the computational analysis contacts displaying highly non-physiological activity. SEEG signals were preprocessed and analyzed using custom-made code in Python 3 based on the Numpy, Scipy and MNE libaries. Signals were analyzed in the referential montage (reference to CPz). Prior to the main analysis, they were low-pass filtered with a zero-phase FIR filter with cutoff at 700 Hz and stopband at 875 Hz (175 Hz transition bandwidth, −6 dB supression at 787.5, maximal ripples in passband 2%) to remove aliasing effects. A high-pass zero-phase FIR filter with cutoff at 1 Hz and stopband at 0 Hz (1 Hz transition bandwidth, −6 dB supression at 0.5 Hz, maximal ripples in passband 2%) was also applied to remove slow drifts from the SEEG signals. Additionally, we also used a band-stop FIR filter at 50 Hz and its harmonics to remove the power line interference (1 Hz band-stop width, 53 dB attenuation at center frequency, maximal ripples in passband 2%).

In addition, we identified time periods containing widesperad artifacts or interictal epileptic events such as spikes, using the procedure described in (Arnulfo et al., 2015). Following this method, we used Morlet wavelets (width *m* = 7) to obtain a spectral decomposiotion of each signal into 33 logarithmically scaled frequencies from 2 to 512 Hz in steps of 1/4 octave. Then, we divided the signal envelopes in non-overlapping 250 ms temporal windows. Corrupted temporal windows were defined as those where at least 10% of contacts had amplitude envelopes 5 standard deviations above their respective mean amplitude in more than half of the 33 frequency bands.

### 2.6 Event-related potentials

Before conducting the spectral analysis, we analyzed the average signal across trials for a selection of relevant recording sites in each subject. For subject 1, our relevant contact was placed in V1/V2 (deepest electrode contact of the trajectory highlighted in Fig. 1B). To compute the event-related potential (ERP) that was locked to the stimulus-onset and avoid potential noise due the low number of trials, we further low-pass-filtered the data under 30 Hz (FIR filter with 6 dB supression at 33.75 Hz, maximal ripples in passband 2%). We then epoched the data (−500 ms to 1500 ms from stimulus onset), baseline-corrected each epoch (subtraction of the mean across the baseline period, from −500 to 0 ms) and extracted the median across trials.

For subject 2, our relevant contact was placed in the left-hemisphere primary motor cortex (M1), approximately at the location of right-hand movement control. In this case, we extracted the ERP in M1 during recognized and non-recognized trials, time-locked to button press or response timeout end, respectively. To compute the ERPs, we low-pass-filtered the data under 30 Hz, epoched the data (500 ms before stimulus presentation to 500 ms after button press or response timeout end, respectively), baseline-corrected each epoch (subtraction of the mean across the baseline period, from −500 to 0 ms before stimulus presentation), aligned the data to button press or response timeout end, respectively, chunked each epoch from −500 to 500 ms from behavioural response and extracted the median across trials of each type (recognized vs non-recognized). In this case, we tested differences in the signals across both conditions using the following procedure: Experimental conditions were compared using a Ranksum test at each time point, with a criterion of *P* < 0.05 for a minimum of 102 consecutive samples (50 ms).

### 2.7 Spectral estimates

Spectral power was estimated from 4 to 512 Hz using an adaptive multitaper method based on discrete prolate spheroidal sequences (DPSS, aka. Slepian sequences) (Slepian and Pollak, 1961; Thomson, 1982; Mitra and Pesaran, 1999). For our analysis, we used custom-made code to achieve the highest flexibility in adjusting the temporal and frequency smoothing for each frequency independently. Following this approach, we sought to find the best temporal resolution at lower frequencies, while obtaining more accurate power estimates at typically low SNR (signal-to-noise ratio) higher frequencies, at the expense of temporal and frequency resolution.

As suggested by previous literature (Buzsáki and Draguhn, 2004; Hipp et al., 2012), both the mean frequency and bandwidth of meaningful brain activity typically follow a logarithmic progression. Low-frequency activity (theta, alpha, beta) is thought to be oscillatory, frequency-specific, and less spatially localized, reflecting a sum of different contributions, in particular widespread post-synaptic potentials. On the other hand, high frequency activity (gamma, high-gamma and above) has a broadband profile and is typically thought to reflect locally synchronized neuronal activity. However, the specific frequency range of such activity is not well established and can vary depending on the recording technique (Buzsáki and Draguhn, 2004).

To consistently capture the specificities of low and high frequency activity, we computed power estimates across 29 logarithmically scaled frequencies *f* from 4 to 512 Hz in steps of 1/4 octave, i.e., each frequency was obtained by multiplying the previous one by 2^1/4^. In addition, we adjusted the spectral smoothing parameter to match 3/4 octave, yielding a spectral resolution of [*f* – 0.34*f*, *f* + 0.34*f*] for each frequency of interest *f*. Regarding the temporal resolution, we followed a different approach for frequencies above and below 32 Hz. For every frequency of interest *f* below 32 Hz, we adjusted time windows to include 6 cycles of *f*, using shorter windows for larger frequencies. This yielded a temporal smoothing ranging from *t* ± 750 ms at 4 Hz to *t* ± 95 ms at 32 Hz around each time point *t* for our estimates. With this temporal and frequency smoothing, we could use a total of 3 tapers for the spectral estimates at each frequency. In contrast, for frequencies above 32 Hz, we used a fixed temporal smoothing of ±100 ms around each time point (total temporal smoothing of 200 ms), which allowed us to use a greater number of tapers for larger frequencies (from 3 tapers at 32 Hz to 69 tapers at 512 Hz), thus increasing the signal-to-noise ratio of our spectral estimates. Independent power estimates were obtained by projecting the signals onto each taper. Then, single-taper estimates were averaged across tapers, thus obtaining a single power time course for frequency.

To avoid undesired boundary effects, we first obtained spectral estimates across the whole task period and epoched the data afterwards. For stimulus-related responses, we considered the epoch comprised from 500 ms before stimulus presentation to 1500 ms after stimulus presentation (this includes the stimulus period and a post-stimulus period of at least 500 ms). Trial periods containing corrupted time windows as defined in section 2.5 were discarded. For each trial, time-frequency plots were normalized at each scale by the mean power (division by the mean power at that scale) computed during the baseline period (from −400 to −100 ms, to avoid temporal contamination), as done in (Rey et al., 2014). Then, we took the median across all trials to characterize each contact’s response to the stimulus.

On the other hand, for motor-related responses (recognized and non-recognized), we considered the epoch comprised from 500 ms before stimulus presentation to 500 ms after button press or response timeout end, respectively. For each trial, time-frequency plots were normalized at each scale by the mean power (division by the mean power at that scale) computed during the baseline period (from −400 to −100 ms, to avoid temporal contamination). We then aligned the data to button press or response timeout end, respectively, chunked each epoch from −500 to 500 ms from behavioural response and extracted the median across trials of each type (recognized vs non-recognized).

To validate our assumption that multitaper estimation might better capture local high-frequency activations than other temporally-resolved techniques, we repeated the whole analysis using a wavelet approach, and compared the spectral power estimates. Power estimates were obtained between 4 to 512 Hz using the same 29 logarithmically scaled frequencies *f* as before with Morlet wavelets (*m* = 7). We then epoched, normalized and averaged the spectrograms exactly as we did with the multitaper estimates. In addition, we repeated the same analysis filtering the wavelet power estimates below 8 Hz before epoching and averaging, to obtain a similar temporal smoothing than with the continuous multitaper method. See Supplementary Information and Fig. S1 for results and discussion.

#### 2.7.1 Statistical inference of task-related activations

When assessing stimulus-related activations, the median of the spectrograms obtained by means of multitaper power estimation was computed over all face presentation trials. Following (Rey et al., 2014), we visually identified time-frequency windows of interest (TFOIs) related to the stimulus presentation on the trial-median spectrograms of certain recording sites of interest (for instance, marked with black rectangles in Fig. 2B). We then quantified the strength in the LFP power response in the defined TFOIs using the following non-parametric method. For each TFOI we defined a surrogate baseline TFOI (window spanning the same frequency ranges and time-width of the TFOI centered in the baseline period). For each trial, we extracted the mean power within the TFOI and the surrogate TFOI. We used a Ranksum test to evaluate significance of TFOI mean power against its surrogate baseline.

**Figure 2:**
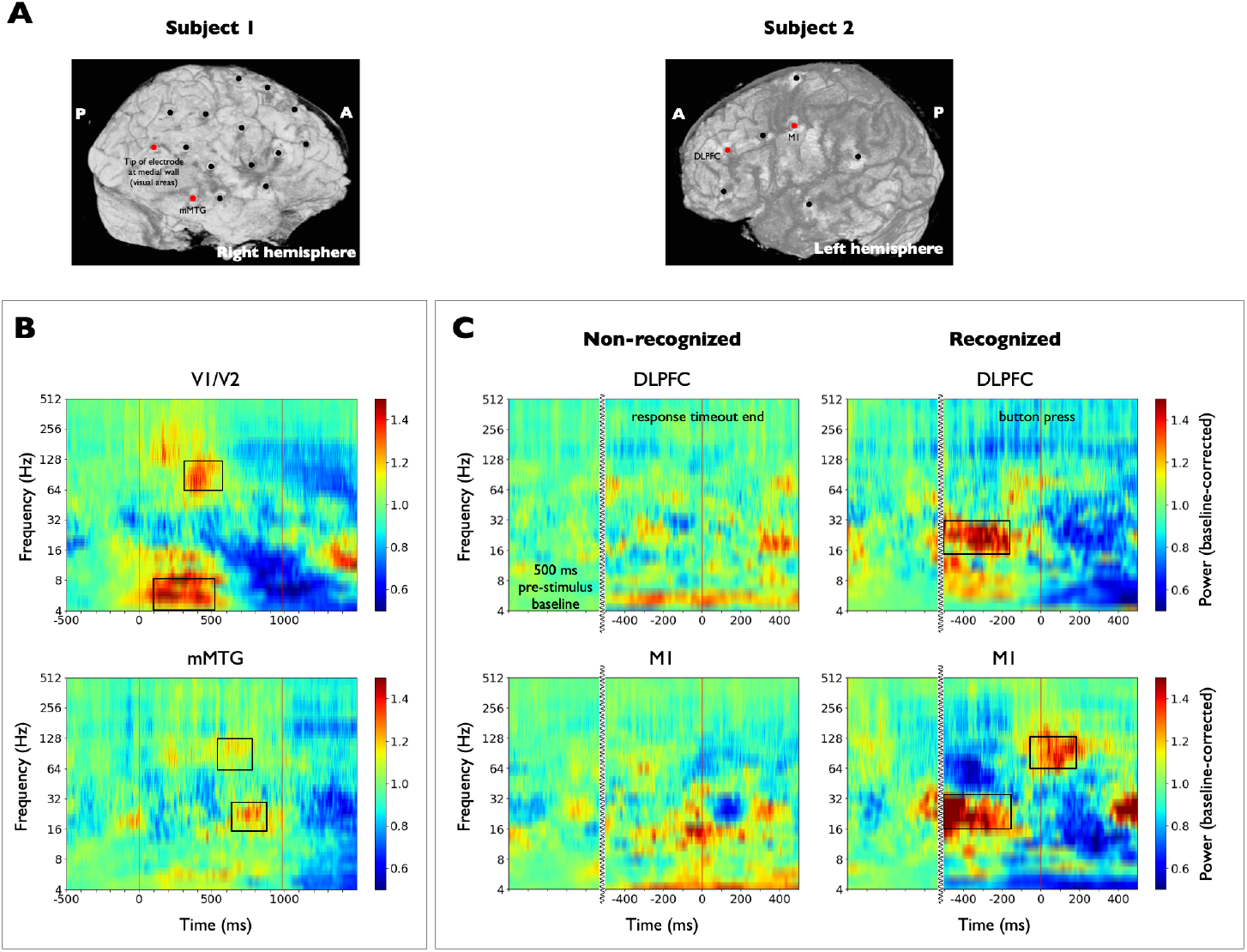
Spectral activations in different task-relevant ROIs for each subject. Spectral activations in different task-relevant ROIs for each subject. **(A)** 3D brain reconstructions showing the electrode entry points for subjects 1 and 2 (black dots). The full trajectory and end point of the electrode targeting V1/V2 in subject 1 can be seen in Fig. 1B. Red dots highlight electrodes for which spectrograms are shown in (B) and (C). **(B)** Median across all trials of the baseline-corrected spectrograms aligned to stimulus presentation (0 ms) and obtained from two different key recording sites in the visual stimulus processing pathway of subject 1. The time-frequency windows of interest (TFOIs) used for statistical comparisons are marked with black rectangles. There is an early power increase in the theta band (*P* < 10^-5^) in the visual cortex (V1/V2, same contact as in Fig. 1C) followed by high-gamma discharges (*P* < 10^-5^). In the mMTG (middle portion in the anterior-posterior axis of the middle temporal gyrus) we found power activations during the second half of the stimulus presentation both in the beta (*P* < 0.001) and the high-gamma range (*P* < 0.001). Vertical red lines indicate the stimulus onset and offset times, respectively. **(C)** Median baseline-corrected spectrogram of the single-trial LFP power across recognized and non-recognized trials aligned to button press or response timeout end (0 ms, red vertical line), respectively, obtained from two different areas related to perceptual decision making and motor report. Power increases in the beta band until 250 ms before button press were found to be significant both in DLPFC (dorsolateral prefrontal cortex) and M1 (same contact as in Fig. 1D) with respect to baseline distribution (*P* < 0.01), but not significant with respect to non-recognized trials. A significant increase in high-gamma power was found in M1 around button press both with respect to baseline (*P* < 0.01) and to the non-recognized trials (*P* < 0.01). Pre-stimulus baseline is also shown for comparison. Curvy lines mark a discontinuity in time.

When assessing motor-related activations locked to the behavioural response (only subject 2), we computed the median of the spectrograms across the two conditions (recognized vs non-recognized) separately. In recognized trial-median spectrograms we visually identified time-frequency windows of interest (TFOIs) and tested the significance of their activations against the baseline period and against the non-recognized trials independently. Significance with respect to baseline was tested using the procedure described in the previous paragraph. Significance with respect to non-recognized trials was tested using a Ranksum test on TFOI mean power values between conditions.

### 2.8 Global connectivity variables

To characterize the global connectivity state, we used two independent and complementary connectivity measures commonly used in the iEEG literature to quantify statistical relationships between signals: the functional connectivity (FC; Cruzat et al., 2018; Tauste Campo et al., 2018) and the phase-locking value (PLV; Axmacher et al., 2008; Arnulfo et al., 2015). On one hand, the functional connectivity (FC) is a linear measure based on amplitude correlations. It is computed as the Pearson correlation coefficient of signal time courses across time. On the other hand, the phase-locking value is a non-linear measure based exclusively on phase couplings. It quantifies the consistency of phase differences between signal time courses across time. For the purpose of connectivity computation, we considered all pairs of GM contacts, except those that were simultaneously in the same electrode and within the same ROI, since they typically measure very similar activity. Note that in SEEG, the electrodes are placed only in some brain areas, which might vary from subject to subject. The mapping of neuronal connectivity is therefore limited to these areas, in contrast to connectivity across all cortical regions, as in fMRI or MEG.

#### 2.8.1 Time-resolved mean functional connectivity

We estimated the time-resolved functional connectivity (FC) between recording sites (broadband signals) using a sliding-window approach with a window length of 200 ms (410 samples) and a step of 1 sample. Before computing the FC, we high-pass filtered the signals above 5 Hz to have at least one full cycle within the 200-ms window (FIR filter with 6 dB supression at 4 Hz, maximal ripples in passband 2%)). When assessing the stimulus-locked global fluctuations, we aligned the signals’ time courses to stimulus presentation, epoched the data from −500 to 1500 ms around stimulus onset and estimated the time-varying FC for a pair of signals using the following procedure. First, we computed the Pearson correlation coefficient between both signals in each time window and trial. For each time window, the FC between two contacts *k*_1_, *k*_2_ was computed as the magnitude of the average correlation value across trials:

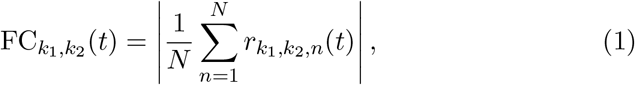

where *N* is the number of trials and *r*_*k*1_, *k*_2,*n*_(*t*) represents the Pearson correlation coefficient between signals *k*_1_ and *k*_2_ in the time window starting at time *t* of the *n*-th trial.

To summarize pairwise connectivity values into a single network metric, we defined the time-resolved mean FC (mFC) as the mean network strength, i.e., the average of FC values over all connections (excluding pairs of contacts simultaneously in the same electrode and within the same ROI). Correlation values were Fisher’s z transformed before taking averages across connections. With this procedure, we obtained a single mFC value for each time window. For each subject, we visually identified one time window with a significant deflection of the mFC with respect to the baseline period and tested for significant differences using a Ranksum test across trials on the time average mFC in the baseline period and in the selected window, with a criterion of *P* < 0.05.

On the other hand, when assessing the motor-related global fluctuations, we selected recognized trials, aligned the signal’s time courses to button press, epoched the data from −500 to 500 ms from button press and estimated the time-varying mFC using the procedure described before. We generated a surrogate mFC time-course using the non-recognized trials. In this case, we aligned the signal’s time courses to response timeout end, epoched the data from −500 to 500 ms from response timeout and estimated the time-varying mFC.

The main analysis was performed by computing the broadband mFC (Tauste Campo et al., 2018), i.e., by estimating the Pearson correlation between broadband signals. Fluctuations captured by this measure might be explained by frequency-dependent correlations. To account for such effects, we also computed the mFC in the alpha (8 – 12 Hz), beta (12 – 30 Hz), gamma (30 – 70 Hz) and high-gamma (70 – 150 Hz) bands independently (See Fig. S2). To do so, we computed the Pearson correlation between band-filtered signals using the same procedure described above.

#### 2.8.2 Time-resolved mean phase locking value

We estimated the time-resolved phase-locking value (PLV) between recording sites across 29 logarithmically scaled frequencies *f* from 4 to 512 Hz using the multitaper method with the same parameters and procedure specified in section 2.7. For every frequency f below 32 Hz, we used a window length of *T* = 6/*f* (6 cycles of *f*) and a step of 1 sample. For frequencies above 32 Hz, we used a window length of 200 ms and a window step of 1 sample. First, analytic signal estimates were obtained independently for each taper, frequency, and contact. Then, single-taper estimates were averaged across tapers and phase time courses were extracted from the taper-averaged analytic signals using the Hilbert transform. When assessing the stimulus-locked global fluctuations, phase signals were aligned to stimulus presentation and epoched from −500 to 1500 ms around stimulus onset. For each time window, the PLV between two contacts *k*_1_, *k*_2_ was computed as the magnitude of the average complex phase difference across trials (Lachaux et al., 1999):

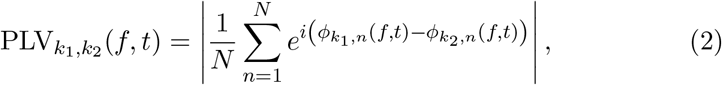

where *N* is the number of trials and *ϕ_k,n_*(*f,t*) represents the phase estimate of contact *k* at frequency *f* and in the time window starting at time *t* of the *n*-th trial. PLV ranges from 0 to 1, where 0 represents maximal phase variability and 1 represents perfect phase locking between signals.

Finally, the mean PLV (mPLV) was defined as the mean network strength, i.e., the average of PLV values over all connections (excluding pairs of contacts simultaneously in the same electrode and within the same ROI). The mPLV was obtained for each time window and frequency, thus obtaining a frequency- and time-resolved global network metric.

To study motor-locked global responses, we computed the mPLV across the two sets of trials (recognized vs non-recognized), independently, as done for the mFC. In this case, phase signals were aligned to button press or response timeout end and epoched from −500 to 500 ms from button press or response timeout end, respectively. We then estimated the time-varying mPLV using the same procedure described above.

### 2.9 Coupling between local activity and global connectivity fluctuations

To study local-global relationships, we aimed to have one time-varying variable *L_k_*(*t*) for each recording site *k* to capture local dynamics (derived from the spectrograms) and one global time-varying variable *G*(*t*) capturing fluctuations in connectivity among recording sites (derived from either the mFC or the mPLV). Based on the preliminary visual inspection, we refined our definition of local activity and defined it as high-gamma activity (64 – 256 Hz) and we analyzed the stimulus-locked setting (subjects 1 and 2). Hence, for each site, the local variable was defined as the average of its (median) spectrogram within the aforementioned high-frequency range. In contrast, at the global level, we used the mFC as defined in the previous section. We also defined a second global variable based on the the mPLV within the frequency range where it showed the highest stimulus modulation (6-16 Hz). To do so, we averaged this variable across the frequency range 6-16 Hz.

To assess the degree of correlation between each contact’s local activity *L_k_*(*t*) and the global connectivity fluctuations measured by the proposed global variable *G*(*t*), we used the following procedure. For each recording site *k*, we computed the Spearman correlation coefficient between its local variable and the global variable across the entire epoch *ρ_k_* = *ρ*(*L_k_*(*t*), *G*(*t*)), where *ρ*(·) stands for the Spearman correlation operator. To infer the significance of each estimation, we built the corresponding null distribution using circular shifts of the local variable, which preserved the local and global variables’ autocorrelation properties (around 200 ms by definition) while destroying their temporal alignment. A detailed description of this methodology can be found in the Supplementary Information. Only medium and large effect size correlations (*r* > 0.3) were kept. In addition, significance level was set to *α* = 0.05 and corrected for multiple comparisons using Bonferroni correction (number of contacts).

A similar procedure was used to assess local-global relationships in the motor-locked setting. In this case, local and global variables were aligned to button press, chunked in temporal windows from −500 to 500 ms, and and only obtained for recognized trials. We then assessed the degree of correlation between each recording site’s local activity and the global connectivity fluctuations using the procedure described in the previous paragraph for the recognized trials.

### 2.10 Data and code availability

Due to institutional restrictions, the data that supports the findings of this study can be accessed only with a data sharing agreement. All code used in this work can be found at https://github.com/mvilavidal/localglobal2022.

## 3 Results

### 3.1 Implantation, behavioural task and basic ERP analysis

We applied our method to the intracranial EEG signals from subjects 1 and 2 while they performed a face-recognition task (Fig. 1A). In this task, each trial consisted of a short pre-stimulus baseline period, an image presentation period and a post-stimulus period in which the subject was instructed to respond within a maximum allowed time.

Subject 1 had 16 electrodes implanted, accounting for a total of 175 contacts, among which 47 were selected for further analysis (see Table 1 and Fig. 1B). Electrodes were implanted on the left hemisphere and covered the primary or secondary visual cortex (1 electrode in V1/V2), the supramarginal gyrus (2 electrodes), large portions of the lateral aspect of the temporal lobe (5 electrodes in the anterior and posterior regions of the STG, MTG, and ITG), anterior and posterior parts of the hippocampus (2 electrodes), the cingulate cortex (4 electrodes distributed uniformly from its rostral anterior to its posterior aspect close to the isthmus), and to a lesser extent the frontal lobe (1 electrode in M1, 1 in the lateral aspect of the PMC and 2 in the DLPFC). For connectivity analyses, we considered all pairs of GM contacts, except those located in the same electrode and within the same ROI, accounting for a total of 1031 pairs. Subject 1 completed N = 64 trials that lasted approximately 12 minutes. Around 0.2% of inspected time windows were marked as corrupt (7/2880). However, none of them affected analyzed periods (from −500 ms to 1500 s around stimulus onset) and no trials were discarded. ERP in the visual cortex of subject 1 was found to be modulated by the stimulus presentation (Fig. 1C), showing negative peaks of –40 μV at around 150 ms after stimulus presentation, with a progressive increase to 50 μV at 500 ms after stimulus presentation.

Subject 2 had 7 electrodes implanted, with a total of 93 contacts, among which 26 were selected for further analysis (see Table 1). Implanted electrodes covered large portions of the frontal lobe (1 electrode in M1, 1 in the medial aspect of the PMC, 2 in the DLPFC and 1 in the VLPFC), the cingulate cortex (5 electrodes distributed from its anterior to its posterior aspect), the STG (2 electrodes) and the insula (1 electrode). For connectivity analyses, we considered all pairs of GM contacts, except those located in the same electrode and within the same ROI, accounting for a total of 307 pairs. Subject 2 completed *N* = 71 trials that lasted approximately 5 minutes. Around 0.3% of inspected time windows were marked as corrupt (3/1200). As before, none of them affected analyzed periods (from −500 ms to 1500 s around stimulus onset; from −500 ms to 500 ms around button press or response timeout) and no trials were discarded. Among all pictures, 22 were reported as recognized, while 49 were reported as non-recognized. In subject 2, event-related potential (ERP) in right-hand M1 (Fig. 1D) displayed significant differences between conditions from −273 to −214 ms before button press and from 4 to 189 ms after button press (Ranksum test at each time point and across conditions, with a criterion of *P* < 0.05 for a minimum of 102 consecutive samples, 50 ms).

### 3.2 Detection of task-driven local activity

The spectrogram of each signal was obtained using the multitaper method. Power estimates at each frequency were then epoched and expressed as a fraction of the mean power during the baseline period (from −400 to −100 ms from stimulus onset) at each frequency, for each trial separately (see Methods, section 2.7).

Subject 1 had an implantation that largely covered the stimulus processing pathway (Fig. 2A, left panel): V1/V2, ITG, MTG, hippocampus. In this case, we studied stimulus-related activations across all trials and disregarded behavioural responses (see Methods, section 2.2). For every contact, the median of the spectrograms was computed over all face presentation trials. Fig. 2B shows the spectrograms of two contacts of subject 1 monitoring V1/V2 and the middle portion (anterior-posterior axis) of the MTG (mMTG) that displayed significant power activations. In V1/V2, we found that the brain responses to the presentation of the stimulus were characterized by an increase in the theta band immediately after the stimulus onset time (100 – 500 ms, 4 – 8 Hz, *P* < 10^-5^) and a high-gamma increase after 300 ms (300 – 600 ms, 64 – 128 Hz, *P* < 10^-5^). In addition, the mMTG exhibited a significant deflection localized in the beta band during the second half of the stimulus period (16 – 32 Hz, 600 – 900 ms, *P* < 0.001), accompanied by a more diffuse activation in the high-gamma range (64 – 128 Hz, 500 – 800 Hz, *P* < 0.001).

In subject 2, no ROI directly related to visual stimulus processing was monitored, while various areas related to perceptual decision making and motor report were covered (Fig. 2A, right panel): M1 at the position of the right-hand (used for button pressing), premotor areas, VLPFC and DLPFC. We first performed the stimulus-related analysis. Here, we also studied motor-related activations in recognized trials. For every recording site, the median of the spectrograms was computed over each set of trials aligned to button press or response timeout end, respectively (see. Fig. 2C). In this setting, we found significant power activations in the beta band of the the DLPFC before button press (Fig. 2C, 16-32 Hz, from 500 to 250 ms before button press, *P* < 0.01). These power activations were, nonetheless, nonsignificant when compared to non-recognized trials. Significant beta power activations were also found in the same time-frequency band in the primary motor cortex (16-32 Hz, from 500 to 250 ms before button press, *P* < 0.01). These were also non-significant when compared to non-recognized trials. In addition, M1 displayed high-gamma activations around button press (64-128 Hz, from 50 ms before to 200 ms after button press). Unlike the previous cases, the activations here were significant when compared both to the baseline period (*P* < 0.01) and to the non-recognized trials (*P* < 0.01).

Overall, the detected ROIs at early and late stages of the cortical face recognition and report pathway, i.e. V1/V2 and M1, provided statistical evidence of activations at frequency ranges (Buzsáki et al., 2012) and times (Salinas and Romo, 1998; Van Vugt et al., 2018) that were compatible with local activity encoding visual stimulus and motor report information. Based on these results, we chose to define the local variables of the local-global analysis within the frequency range 64 – 256 Hz.

### 3.3 Global connectivity effects of stimulus presentation and motor response

The mean functional connectivity (mFC) and the mean phase locking value (mPLV) were computed as described in Methods (see section 2.8). The upper panel of Fig. 3 shows the time evolution of both variables in the stimulus-locked setting for each subjet (Fig. 3A, mFC; Fig. 3B, mPLV). Despite having different implantation schemes, a decrease in the mFC associated to stimulus presentation was consistently found in both subjects with respect to their pre-stimulus baseline distribution in a temporal window of 300-600 ms. To test significance, a Ranksum test was applied across contact pairs on the average FC in the baseline period and the 300-600 ms window (*P* < 10^-6^, Cohen’s D effect size *D* > 5 in both subjects, Fig. 3A). Figure S2 shows the time evolution of mFC when computed in the physiological bands. The broadband mFC decrase was found to be localized in the alpha (8 – 12 Hz) and, to some extent, beta bands (12 – 30 Hz). This decrease in time-resolved mFC (time windows of 200 ms) was accompanied by a concurrent decrease in mPLV in the frequency range 6-16 Hz that lasted 200 ms (z-score < –3, Fig. 3B).

**Figure 3:**
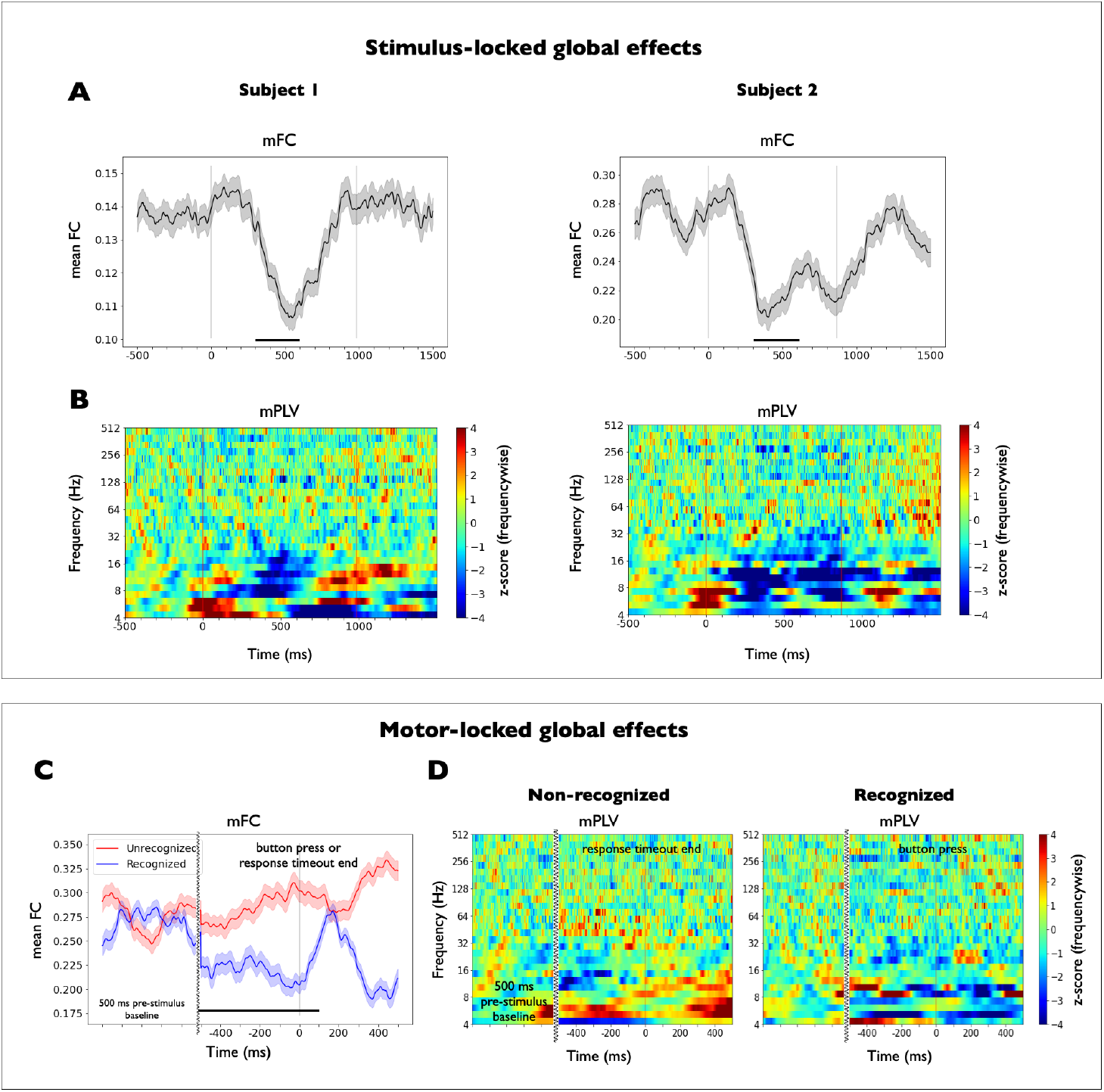
Stimulus- and motor-related global effects. Stimulus- and motor-related global effects. The upper panel shows the temporal evolution of the global variables aligned to stimulus presentation in subjects 1 and 2. **(A)** Time course of the broadband mean functional connectivity (mFC) aligned to stimulus onset. Mean ± SEM across all GM contact pairs (*N*_pairs_ = 1031 in subject 1, *N*_pairs_ = 307 in subject 2). Correlation values were Fisher’s z transformed before taking averages across contacts pairs. See Methods, section 2.8, for details. Vertical dark lines indicate the stimulus onset and offset times, respectively. A significant decrease in the mFC associated to stimulus presentation was found in the two subjects with respect to their pre-stimulus mean mFC value (300-600 ms, marked with black bars, Ranksum test applied across contact pairs on the average FC in the baseline period and the 300-600 ms window, *P* < 10^-6^, Cohen’s D effect size *D* > 5 in both subjects). **(B)** Time-resolved mean phase locking value (mPLV) aligned to stimulus presentation. See Method, section 2.8, for details. Plots have been z-scored with respect to the pre-stimulus period (from −500 to −100 ms) at each frequency scale for visualization purposes. Vertical red lines indicate stimulus onset and offset times, respectively. A significant phase-decoupling was consistently observed across trials in the frequency range 6-16 Hz around 300 ms after stimulus presentation in the two subjects (z-score < 3). This effect lasted around 200 ms. The lower panel shows the evolution of the global variables aligned to the behavioural response across the corresponding set of trials in subject 2. **(C)** Time course of the broadband mean functional connectivity (mFC) for subject 2 in recognized (N=22) and non-recognized (N=49) trials. Mean ± SEM across all GM contact pairs (*N*_pairs_ = 307) for each set of trials aligned to button press or response timeout end (marked with a vertical dark line), respectively. Pre-stimulus baseline is also shown for comparison. Curvy lines mark a discontinuity in time. A significant mFC difference between conditions was found between 500 ms before and 100 ms after button press or response timeout end, respectively (marked with a black bar, Ranksum test applied across contact pairs on the average FC in the selected time window across conditions, *P* < 10^-6^, Cohen’s D effect size *D* > 5). **(D)** Time-resolved mean phase locking value (mPLV) for subject 2 in recognized (N=22) and non-recognized (N=49) trials, aligned to button press or response timeout end, respectively. Plots have been z-scored with respect to the pre-stimulus period (from −500 to −100 ms before stimulus presentation) at each frequency scale for visualization purposes. Vertical red lines indicate button press or response timeout end, respectively. Pre-stimulus baseline is also shown for comparison. Curvy lines mark a discontinuity in time. No significant differences could be found between both conditions in the frequency range 6-16 Hz around button press / response timeout end.

To show that the reported connectivity effect was of widespread nature we performed two control analysis. First, we inspected the influence of power fluctuations into the mFC and mPLV decreases. For mFC, we plotted the time-resolved average standard deviation and covariance across all contact pairs in the same period (See Fig. S3) paying special attention at the 300-600 ms window after stimulus onset. The curves illustrate that a prominent decrease was manifested for both subjects in their average covariance (Fig. S3B) while only mild increases were observed in the average standard deviation, i.e., in the power amplitude of the signals (Fig. S3C), suggesting that the observed mFC decay did not result from mere power fluctuations and reflected a real decrease in connectivity. For mPLV, we assessed whether the decrease in 6-16 Hz was due to phase decoupling or simply reflected a decrease in power. To do so, we compared the average time/frequency power with the mPLV (Fig. S4). No global decrease in energy was observed. Hence, the decay in mPLV could not be explained by a decrease in the signal-to-noise ratio or a simple decrease in power in 6-16 Hz. This suggests that a genuine phase decoupling in this frequency range. Second, we assessed how the connectivity decay in both variables varied across all contacts (See Fig. S5 and Fig. S6). Indeed, Fig. S5B and Fig. S6B suggest that the period 300-600 ms was affecting the connectivity strength of a substantial subset of recording contacts (see marked rectangles) while the curves for pre-stimulus and stimulus epochs displayed in Fig. S5C and Fig. S6C confirmed that the decay was a generalized effect across the implantation scheme of both subjects. In addition, the connectivity matrices shown in Fig. S7 highlight that the decay is distributed over a great proportion of contact pairs.

With regard to the motor-locked setting, the lower panel of Fig. 3 shows the time evolution of the two global variables for subject 2. In this setting, trials were aligned to button press or response timeout end, and global variables were computed in a time period from −500 to 500 ms across recognized (N=22) and non-recognized (N=49) trials, respectively (Fig. 3C, mFC; Fig. 3D, mPLV). A significant mFC difference between conditions was found between 500 ms before and 100 ms after button press or response timeout end, respectively (Ranksum test applied across contact pairs on the average FC in the selected time window and across conditions, *P* < 10^-6^, Cohen’s D effect size *D* > 5). Here, no significant differences with respect to the pre-stimulus baseline or across conditions were found with mPLV in the frequency range 6-16 Hz.

### 3.4 Local-global relationships

We here investigated the relationship between the local variable of each recording site (average power across the high-gamma range, 64-256 Hz) and the two global connectivity-based variables (mFC, and mPLV averaged across the range 6-16 Hz). This relationship was explored in the two proposed settings (stimulus-locked, motor-locked) to find potential local-global couplings associated to stimulus information processing and motor reports (see Methods, section 2.9)

#### 3.4.1 Stimulus-locked local-global coupling

Fig. 4 shows the results for the stimulus-locked local-global analysis in subjects 1 (Fig. 4A and 4B) and 2 (Fig. 4C and 4D). In particular, the upper left plot in Fig. 4A shows the time evolution of each contact’s local variable (average power across the high-gamma range, 64-256 Hz) across the stimulus presentation in subject 1. Contacts were sorted by the mean high-gamma power in the 300–600 ms time window after stimulus presentation, which defines the task-related activation of each recording site (shown next to each contact’s power evolution). The lower plot in Fig. 4A shows the time evolution of the two global variables (mFC, and mPLV averaged across the range 6-16 Hz) temporally aligned to the local variables. The results of local-global testing are shown in Fig. 4B for each global variable, separately. To assess the degree of local-global correlation we computed the Spearman correlation coefficient between each local variable and the global variable across the entire epoch. Only medium and large effect-size correlations (*r* > 0.3) were considered. Significance level was set to α = 0.05 (null distribution built using circular shifts of the local variable) and corrected for multiple comparisons across multiple contacts. In subject 1, the local increase of activity of high-gamma activity at V1/V2 was found to have a significant correlation of −0.7 with the decrease in the two global variables (see examples of downsampled scatter plots for each global variable in S8A). The decrease of mFC was also significantly correlated with high-gamma activations in other contacts such as mMTG (middle portions of the MTG, *r* ≈ –0.5), which also ranked high in task-related activations. The fluctuations in mPLV correlated also with other contacts that were less active during the task. A significant negative correlation was found with the PCC. In addition, some contacts that suffered negative deflections in high-gamma activity correlated positively with mPLV fluctuations (PMC, posterior portion of the STG, *r* ≈ 0.5).

**Figure 4:**
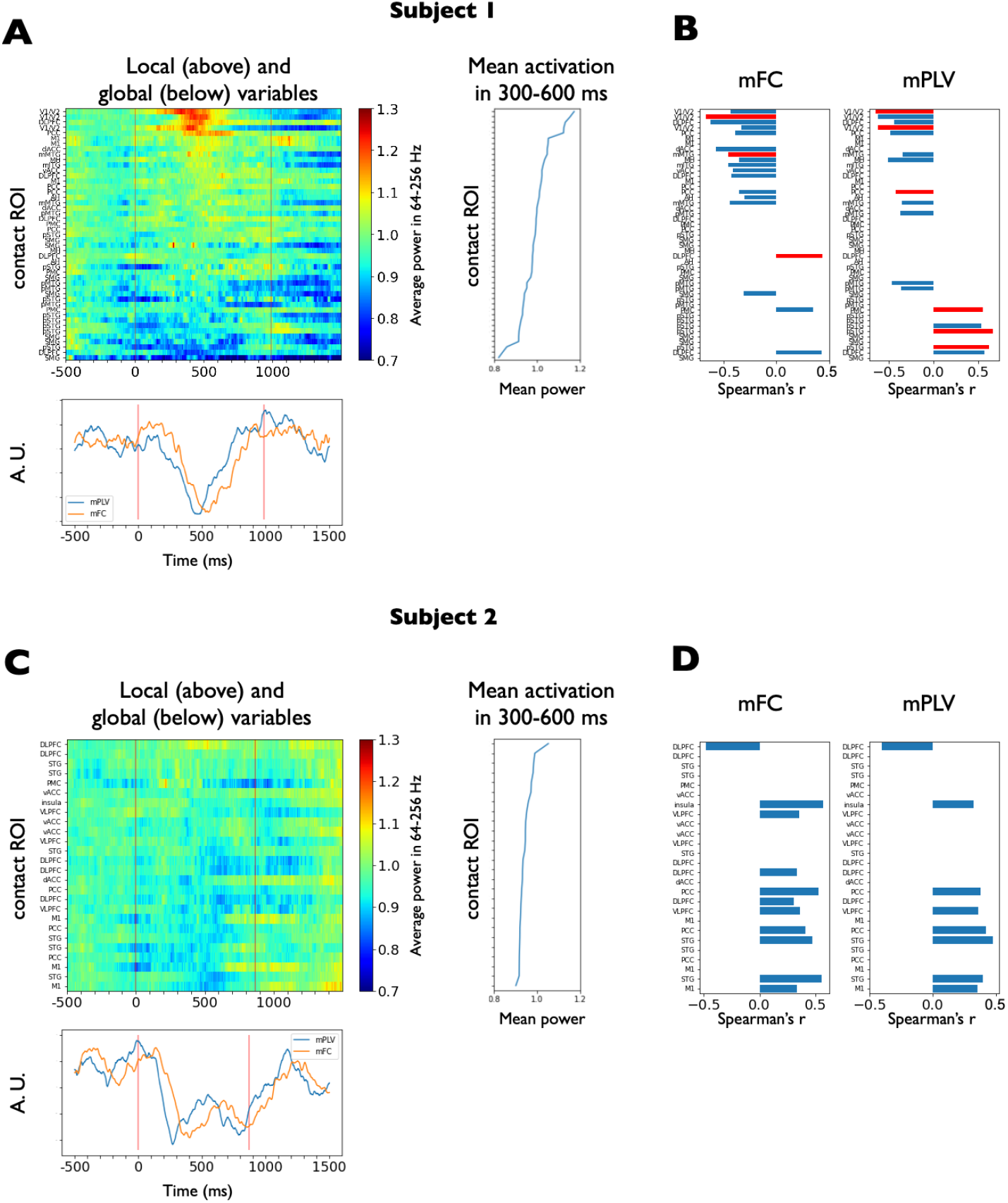
Stimulus-related local-global coupling. Stimulus-locked local-global coupling for subjects 1 (upper panel: subfigures A and B) and 2 (lower pannel: subfigures C and D). **(A)** Local and global variables for subject 1. (Upper) Local variable for each brain recording site in subject 1 (median spectrogram power across all trials, averaged across the high-gamma range, 64-256 Hz, see Methods, section 2.9). Vertical red lines indicate the stimulus onset and offset times, respectively. Contacts have been sorted by the mean high-gamma power in the 300–600 ms time window after stimulus presentation, which defines the task-related activation of each contact (shown next to each contact’s power evolution). (Lower) Global variables: mFC, and mPLV averaged within the frequency range 6-16 Hz. Global variables have been rescaled for the purpose of comparison (shown in arbitrary units, A. U.). Vertical red lines indicate the stimulus onset and offset times, respectively. **(B)** Spearman correlation coefficient between each local variable and each global variable in subject 1. Correlations have been thresholded at medium-size effects (*r* > 0.3). In addition, significance was tested using a surrogate distribution via circular shifts, with a significance criterion of *α* = 0.05 and corrected for multiple comparisons for the number of contacts. Significant correlations are indicated with red bars. Local increase in activity at V1/V2 appears to have a significant correlation of −0.7 with the decrease in the two global variables. In addition, The decrease in mFC seems to have a significant correlation with local activity in task-relevant contacts such as mMTG and mITG (middle portions of the MTG and ITG, respectively). The local activity at PCC appears to be negatively correlated with mPLV. **(C)** Local and global variables for subject 2. Analogous to (A) **(D)** Spearman correlation coefficient between each local variable and each global variable for subject 2. Analogous to (B). However, in this case, none of the local variables showed a significant correlation with the global variables.

In an analogous form, the upper left plot in Fig. 4C shows the time evolution of the local variable (average power across the high-gamma range, 64-256 Hz) across the stimulus presentation in subject 2. High-gamma activity appears to be much less modulated by the task than in subject 1 (see task-related activation of each recording site next to each contact’s power evolution), probably due to an implantation scheme covering regions that were not involved in the initial processing phases of visual stimuli. The lower plot in Fig. 4C shows the time evolution of the two global variables (mFC, and mPLV averaged across the range 6-16 Hz) temporally aligned to the local variables. The results of local-global testing are shown in Fig. 4D for each global variable, separately. Interestingly, although global fluctuations in subject 2 exhibited very similar trends to those observed in subject 1, no local variable significantly explained the reported fluctuations.

#### 3.4.2 Motor-locked local-global coupling

We here tested couplings between the global variables and the local ones when aligned to button press (time period from −500 to 500 ms from motor report) in subject 2. Fig. 5 shows the results for motor-locked local-global analysis in recognized trials. The upper left plot in Fig. 5A shows the time evolution of each contact’s local variable (average power across the high-gamma range, 64-256 Hz) around the motor report in subject 2. Contacts have been sorted by the mean high-gamma power from −100 to 200 ms from button press, which defines the motor-related activation of each recording site (shown next to each contact’s power evolution). The lower plot in Fig. 5A shows the time evolution of the two global variables (mFC, and mPLV averaged across the range 6-16 Hz) temporally aligned to the local variables. In all figures, pre-stimulus baseline is also shown for comparison although not used for local-global analysis. Local-global correlation was assessed using the same procedure as in the stimulus-locked setting. The results of local-global testing are shown in Fig. 5B for each global variable, separately. Importantly, local increases in activity at M1 were found to be positively correlated (*r* = 0.5, *P* < 0.05 corrected for multiple comparisons across contacts; see an exemplary downsampled scatter plot in Fig. S8B left) with mFC, that also exhibited an increase during motor report. One contact in the vACC that showed prominent negative deflections during the motor report period appeared to have significant negative correlations with mFC. This contact also displayed a significant correlation with mPLV. This evidence supports the global nature of the connectivity measures described above. Taken together, the results for both subjects suggest the existence of a physiological link between the stimulus-driven activation of certain areas and slow global connectivity fluctuations.

**Figure 5:**
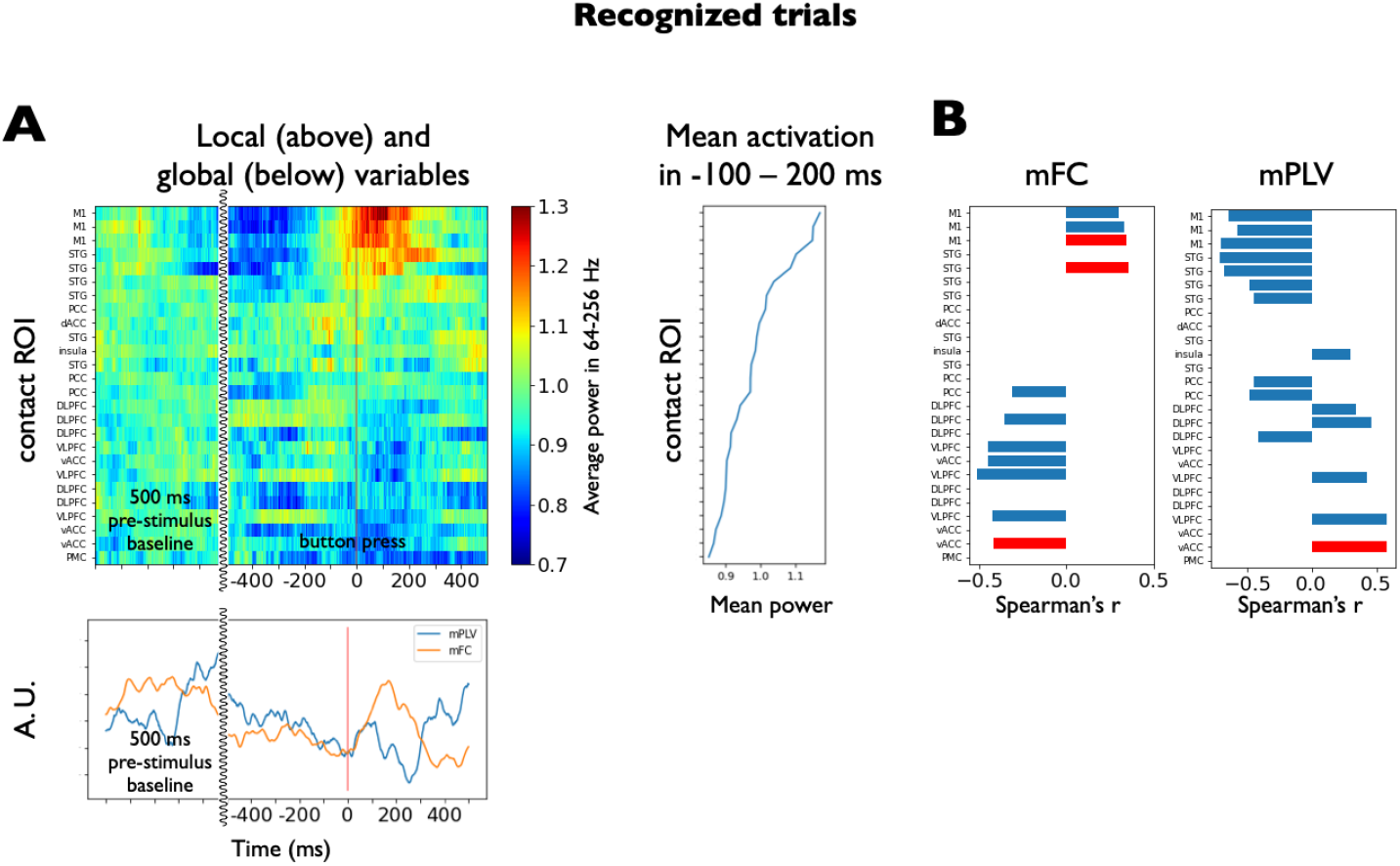
Motor-related local-global coupling for subjects 2 in recognized trials. **(A)** Local and global variables for recognized trials. (Upper) Local variable for each brain recording site in subject 2 aligned to motor report (median spectrogram power across recognized trials aligned to button press, averaged across the high-gamma range, 64-256 Hz, see Methods, section 2.9). The vertical red line indicates button press. Contacts have been sorted by the mean high-gamma power in a time window around button press (from −100 to 200 ms), which defines the motor-related activation of each contact (shown next to each contact’s power evolution). (Lower) Global variables: mFC, and mPLV averaged within the frequency range 6-16 Hz. Global variables have been rescaled for the purpose of comparison (shown in arbitrary units, A. U.). The vertical red line indicates button press. Pre-stimulus baseline is also shown for comparison (not used for correlations). Curvy lines mark a discontinuity in time. **(B)** Spearman correlation coefficient between each local variable and each global variable in the recognized trials during the time period of −500 to 500 ms around button press. Correlations have been thresholded at medium-size effects (*r* > 0.3). Significance was tested using a surrogate distribution via circular shifts, with a significance criterion of *α* = 0.05 and corrected for multiple comparisons for the number of contacts. Significant correlations are indicated with red bars. Local increase in activity in one contact located in M1 has a significant positive correlation of 0.5 with mFC. An additional contact in the STG appears to have a significant positive correlation with mFC. A contact in the vACC shows significant correlations both with mFC and mPLV, but its power fluctuations in the high-gamma range are much lower than those of M1.

### 3.5 Additional analysis

We cross-validated the methodology in a third subject that performed a cognitive task under a slightly different paradigm (see Methods, section 2.2) and compared the outcomes of the analysis with our previous findings. First, we observed neural activations in the gamma and high-gamma bands around 300 ms after stimulus presentation in relevant contacts localized in the vicinity of V1/V2 (see Fig. S9A left, 300 – 600 ms, 64 – 128 Hz, *P* < 10^-^3) similarly to subject 1 (Fig. 2A). In addition, high-gamma activity was detected in the anterior hippocampus starting 500 ms after stimulus presentation (Fig. S9A right, 500 – 1000 ms, 64 – 128 Hz, *P* < 10^-^4).

In the stimulus-locked setting, a consistent decay in global connectivity was observed after stimulus presentation. Although the mFC exhibited high baseline fluctuations that yielded a non significant output (Fig. S9B left), the connectivity decay present in subjects 1 and 2 (Fig. 3B) was clearly captured by the mPLV as a phase decoupling within the frequency range 6-16 Hz. Accordingly, local-global coupling analysis was performed with mPLV (Fig. S9D and S9E), which was found to be strongly modulated by the task. Indeed, local increases in activity at V1/V2 and STG contacts, respectively, were found to negatively correlate with the decrease in mPLV (r ≈ –0.6). In addition, the activity increase in AH during the second half of the stimulus presentation was coupled with enhanced global synchronization.

This third subject was also analyzed under the motor-locked setting. Specifically, subject 3 performed a total of 96 trials, from which 45 were reported as recognized and 51 as not recognized. Concordant with subject 2 (Fig. 3C), a mFC difference between conditions was found between 500 ms before and 100 ms after button press or response timeout end, (Fig. S9C left panel, Ranksum test applied across contact pairs on the average FC in the selected time window across conditions, *P* < 0.05, Cohen’s *D* effect size *D* > 2). An increase in mFC after button press concordant with subject 2 was also found. In a similar vein, the mPLV signatures of recognized and non-recognized trials up to 16Hz (Fig. S9C middle and right panels) also resembled those from subject 2, exhibiting a slight upper shift in the frequency range (See Fig. S9C). Despite the lack of specific (i.e., contralateral to the moved hand) motor-related areas here, both global variables were found to correlate with the local activity of multiple potentially task-engaged areas, such as left MTG, ITG, insula and temporal pole.

## 4 Discussion

In this study we developed and systematized a methodological pipeline integrating neuroanatomic information, signal processing functions and connectivity measures to statistically infer potential correlations between the local activity and global connectivity fluctuations. This new local-global frame-work was defined and tested in two patients with drug-resistant epilepsy with iEEG recordings that performed a face-recognition task. Data from a third subject that performed a cognitive task under a slightly different paradigm was further analyzed to cross-validate some of our method’s assumptions and previous findings with an independently generated dataset. With a small sample size, this study has a methodological conception and its main aim is to present a novel analytical framework to assess local-global couplings in iEEG recordings. The reported biological results should be regarded as a preliminary small-sample size application towards the developed concept.

The crucial point of our framework is the appropriate definition of local and global variables, and their accurate estimation. For the local variables, we used power activations in the high-gamma range (64-256 Hz) estimated via the continuous multitaper method, which is designed to reduce bias with respect to true spectral content (see Supplementary Information for a comparison and discussion of multitaper and wavelet estimation techniques). The global variables were defined by means of functional connectivity analysis. Crucially, they were based on lower frequency cofluctuations, which are thought to have a more widespread origin (Pesaran et al., 2018), reflecting more global states. Based on these premises, we made use of two independent functions in the low-frequency range to measure the brain sites’ coupling consistency across time. On one hand, the mean functional connectivity (mFC) is based on temporal correlations between pairs of broadband signals and captures temporal cofluctuations across recording sites. On the other hand, the mean phase-locking value (mPLV) relies on phase estimation and quantifies the consistency of phase differences between signal time courses (at each frequency). Then, local-global relationships were tested via correlation-based tests, although more sophisticated methods could also be used, as discussed in section 4.4.

### 4.1 Implantation and task analysis

Despite performing the same task, subjects 1 and 2 had differentiated implantation schemes, which allowed to test our framework with heterogeneous conditions. Overall, ROIs at both ends of the cortical pathway in visual perception and motor report were covered when considering both subjects together. Implantation of subject 1 (Fig. 1B) covered extensively the occipital and temporal lobes, including crucial areas of the cortical pathway in visual face perception and processing (V1/V2, ITG, MTG, hippocampus) (Bernstein and Yovel, 2015; Jonas et al., 2016; Wang et al., 2016), while implantation of subject 2 (Fig. 2B) was more extense in the PFC, and motor and premotor areas. Primary results were obtained by aligning trials to stimulus presentation. In addition, we explored modulations aligned to motor-response in recognized trials. Subject 2 was particularly interesting for the purpose of motor-locked analysis, since one contact was located in M1 (left hemisphere), approximately in the region controlling right-hand movement, with which motor reports were made. In this contact, ERPs exhibited a significant deflection when compared to the surrogate condition obtained by using non-recognized trials locked to response timeout end.

### 4.2 Task-driven activations

Following (Rey et al., 2014), we defined certain time-frequency windows of interest (TFOI) in the spectrograms of preselected key recording sites and tested whether activations in those windows were significant with respect to the pre-stimulus presentation or across conditions when aligned to motor response (Fig. 2). Following this approach, in subject 1 we observed significant power increases in V1/V2 in the theta and and high-gamma range during the first half of the stimulus, that were followed by more diffuse activations in the MTG in the beta and high-gamma range during the second half of the stimulus period. Similar activations in the vicinity of V1/V2 in gamma and high-gamma ranges were also observed in subject 3. In this case, additional high-gamma activity was observed in the anterior hippocampus during the second half of the stimulus presentation (starting 600 ms after stimulus presentation). In subject 2 we found significant power activations across all trials in the beta band in the DLPFC and M1 during the second half of the stimulus presentation period. When aligned to motor responses, power activations in the high-gamma range emerged around button press in recognized trials for subject 2.

The above results generalize findings from previous literature related to the face perception pathway found with different recording modalities (Zhen et al., 2013; Rey et al., 2014, 2015; Wang et al., 2016; Grill-Spector et al., 2017; Landi and Freiwald, 2017; Dobs et al., 2019) and help reconstruct additional stages of the visual processing pathway needed to consciously recognize face identities and of the motor planning pathway needed to engage in behaviour. Interestingly, both ends of the cortical pathway (V1/V2 and M1) showed broadband high-frequency activity during stimulus presentation and perceptual report which might reflect the presence of neural populations encoding sensory and motor information in each area, respectively (Buzsáki and Draguhn, 2004). In contrast, intermediate nodes of this pathway such as the MTG area, engaged in processing visual face features, and the DLPFC, engaged in decision planning, exhibited prominent activity in the beta band during the second half of the stimulus presentation. In particular, beta activity in M1 across all trials not concurrent to high-gamma activity might reflect afferent potentials that do not result in local activity.

The encoding of visual stimuli in the visual cortex (Kohn and Smith, 2005; Smith and Kohn, 2008; Graf et al., 2011) and the neural correlates of the somatosensory-motor pathway (Salinas and Romo, 1998; Romo et al., 2002; Luna et al., 2005; Thura and Cisek, 2014; Campo et al., 2015) have long been studied in primates. The novelty of our study, however, lies in having been able to generalize previous results in human brain recordings. In addition, global fluctuations potentially reflecting cognitive states have been poorly analyzed due to the difficulty of recording simultaneous single neurons during task performance. Intracranial electroencephalography (iEEG) recordings from the human brain provide an opportunity to study such fluctuations at the global scale thanks to their coverage of brain areas so distant such as V1 and M1.

### 4.3 Common global fluctuations independent of implantation scheme: Preliminary findings

To characterize global connectivity states during the task, we used two complementary variables. On one hand, the mean functional connectivity (mFC) was defined as the average Pearson correlation across contact pairs in 200 ms time windows, thus capturing cofluctuations of the broadband signals (1-700 Hz), which are ultimately dominated by low frequencies. On the other hand, the mean phase-locking value (mPLV) was defined to characterize the consistency of phase differences between signal time courses at each frequency scale. Significant modulations associated to stimulus presentation in this variable were only found in the 6-16 Hz range. We therefore restricted our analyses to this frequency range for this variable.

Remarkably, despite having differentiated implantation schemes, the connectivity functions consistently showed in both subjects a significant global desynchronization occurring a few hundred milliseconds after stimulus onset (Fig. 3, upper panel), which was shown to be of generalized nature and not specifically biased by power fluctuations (See Figs. S3, S4, S5, S6). In line with these findings, a decay in connectivity was also observed after stimulus presentation in subject 3. Although the mFC exhibited very noisy fluctuations (Fig. S9B left) the effect was clearly captured as a phase decoupling in the frequency range 6-16 Hz measured with mPLV (Fig. S9B right). This suggests that low frequencies in the monopolar montage might be useful when aiming to capture global connectivity states that are independent of the implantation scheme, as already implied by previous works (Tauste Campo et al., 2018). The decrease in time-resolved mFC reflects that the stimulus breaks baseline connectivity cofluctuations, owing to a potential specialization or segregation of different subnetworks in processing the incoming information. This decrease was localized in the alpha and beta bands (Fig. S2), consistently with the results found with mPLV. Note that in our study we used different faces in each trial. Further studies should test whether the trial specificity is maintained when using exactly the same stimulus across trials. When studying motor-locked global effects (subject 2, Fig. 3, lower panel), no clear trend was observed in the mPLV, while the mFC exhibited a transient increase after the response time, an observation that was reproduced in subject 3.

Due to the small sample size and diversity of implantation schemes, these observations should be taken as preliminary results towards the developed concept, in particular considering the large inter-subject and interareal variability of oscillatory signals. Further studies should be performed to assess the consistency of reported results with more subjects and with more trials of each kind. We hypothesize that the stimulus presentation might initially trigger a specialization of the whole-brain network (reflected in the decrease of mFC and mPLV) to process the incoming stimulus in a segregated manner, followed by an increase in connectivity (reflected by the positive deflection in mFC around response) needed to integrate information and plan an internal response. Yet, the underlying mechanism behind the reported global fluctuations remains unclear. Further studies should assess what other variables (context, other cognitive processes, pre-stimulus cognitive state) might modulate these fluctuations. In particular, more ecological frameworks could be used (Freiwald et al., 2016), for instance using dynamic stimuli, to assess the extent to which the reported deflections depend on the stimulus features.

### 4.4 Local-global framework

The local-global framework was used to test whether the activity of some recording sites was statistically coupled to the global fluctuations observed across sites. In particular, we found that the effect highly correlated with the local activity of brain areas involved in visual information processing, providing evidence that the global measures might be a novel signature of functional brain activity segregation taking place when a stimulus is processed in a task context. In addition, in the response-locked setting, the increase in mFC was significantly coupled to the local activity of brain areas in the motor cortex.

Here, we propose a first study to quantitatively asses local-global statistical relationships in a task-related context. Here, we do not aim to capture causal effects, but only time-concurrent local-global phenomena that may plausibly reflect a common functional network. This study should pave the way for more sophisticated methods that can assess the role of each node when considering the whole network together. Generalized Linear Models (GLMs), for instance, could serve this purpose. However, a difficulty in pursuing this approach lies in having insufficient statistical power, specially in short trials with a large number of recording sites. In addition, latent non-observed variables should be considered to control for key hubs in the network that are not monitored but might coordinate global fluctuations. As previously mentioned, such improved models should take into account the modulation of the local-global coupling by other contextual variables or previous cognitive states. Further studies should also investigate directionality to assess direct influences of local activity on global fluctuations and viceversa, for instance by identifying particular global brain states that trigger local activity at certain nodes. This could be done by means of directionality measures such as Granger causality or directed information (Tauste Campo, 2020).

Another issue to take into consideration is the different time scales of local and global fluctuations. Physiologically meaningful local events such as HFOs or neuronal encoding can occur on the order of milliseconds (Romo et al., 2002; Arnulfo et al., 2015). At the same time, some studies have found global fluctuations with characteristic timescales of tens of seconds. In particular, resting state networks (RSN) below 0.1 Hz have been identified under the resting state condition both with fMRI and with time-resolved MEG (De Pasquale et al., 2010; Brookes et al., 2011; Hipp et al., 2012; Buckner et al., 2013). Other studies based on computational modelling or statistical inference have found global network states with a lifetime of around 200 ms (Buckner et al., 2013; Deco et al., 2019). This growing evidence suggests that different brain phenomena can be characterized by diverse spatial scales and evolve over the course of different temporal scales (Northoff et al., 2020; Vila-Vidal et al., 2020). Experimental designs and models linking activity at different spatial scales will have to face this phenomenon. Long resting-state iEEG recordings, for instance, could be used to test this hypothesis based on local intrinsic fast activations linked to slower-changing global states.

### 4.5 Study limitations

The two main limitations of this study are the low number of patients and the limited spatial sampling inherent to the SEEG technique. Although this study has a methodological nature, some of the results reported here should be validated with a larger number of subjects, given the large inter-subject variability of intracranial signals. In particular, patients should be chosen carefully according to their implantation schemes to better cover the visualmotor pathway in the aim to refine and better understand the bidirectional coupling between local activity and global fluctuations during this type of task.

In addition, our framework has been used to correlate fluctuations in global and local variables estimated by leveraging both on time and trials simultaneously as done in previous studies (PLV, Arnulfo et al. (2015); FC, Cruzat et al. (2018)). The main advantage of this procedure is that it provides robust estimates when the number of trials is low, but inter- and intra-trial variabilities become intermingled and are impossible to separate. Although out of the scope of this study, our framework could be adapted to regress the variability of global variables across trials at each time point using the local variables. This could be achieved by using single-trial estimates. However, time autocorrelation in low frequencies supposes a major drawback in pursuing this approach, specially in short trials where slow fluctuations cannot be captured. All in all, further studies should be designed with a larger number of trials to have sufficient statistical power to independently quantify inter- and intra-trial variabilities.

## 5 Acknowledgments

We thank all subjects for their participation in the study. We also thank Rodrigo Quian Quiroga and Hernán G. Rey for sharing some of their code to design the task paradigm. MVV was supported by a fellowship from “la Caixa” Foundation, Spain (ID 100010434, fellowship code LCF/BQ/DE17/11600022). MVV and ATC were supported by the Bial Foundation grant 106/18. GD and ATC were supported by the project “Clúster Emergent del Cervell Humà” (CECH, ref. 001-P-001682), within the framework of the European Research Development Fund Operational Program of Catalonia 2014-2020. GD was supported by a Spanish national research project (ref. PID2019-105772GB-I00 MCIU AEI) funded by the Spanish Ministry of Science, Innovation and Universities (MCIU), State Research Agency (AEI); HBP SGA3 Human Brain Project Specific Grant Agreement 3 (grant agreement no. 945539), funded by the EU H2020 FET Flagship programme; SGR Research Support Group support (ref. 2017 SGR 1545), funded by the Catalan Agency for Management of University and Research Grants (AGAUR); Neurotwin Digital twins for model-driven non-invasive electrical brain stimulation (grant agreement ID: 101017716) funded by the EU H2020 FET Proactive programme; euSNN European School of Network Neuroscience (grant agreement ID: 860563) funded by the EU H2020 MSCA-ITN Innovative Training Networks; Brain-Connects: Brain Connectivity during Stroke Recovery and Rehabilitation (id. 201725.33) funded by the Fundacio La Marato TV3; Corticity, FLAG–ERA JTC 2017, (ref. PCI2018-092891) funded by the Spanish Ministry of Science, Innovation and Universities (MCIU), State Research Agency (AEI).

## Supplementary Information

### S1 Spectral power estimation with wavelet and continuous multitaper methods

Although we chose to use multitaper estimation with DPSS as the main technique to obtain our estimates (used in previous studies such as (Hipp et al., 2012; Lopez-Persem et al., 2020)), we decided to compare the performance of this method with the more widely used wavelet transform method (Rey et al., 2014; Arnulfo et al., 2015; Siebenhühner et al., 2020). Spectral estimation on the time-frequency domain is limited by the fundamental Gabor (aka. Gabor-Heisenberg) uncertainty principle 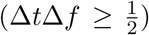, where where Δ*t* and Δ*f* represent the uncertainties in time and frequency (Gabor, 1946). In this context, Morlet wavelets are designed to be almost optimal with respect to the previous inequality, yielding the best possible time-frequency resolution, with Δ*t*Δ*f* ≈ 0.883 (Morlet et al., 1982; Russell and Han, 2016; Cohen, 2019).

On the other hand, continuous multitaper estimation is designed to perform estimates with reduced bias with respect to true spectral content. This is achieved at the expense of temporal and/or frequency resolution. In particular, this method obtains multiple independent estimates of the same sample, projecting the data onto orthogonal functions or tapers (DPSS, in our case). The maximum number of independent tapers (*K*) is given by *K* = Δ*t*Δ*f* – 1. In our case, we adapted the parameters to increase the number of tapers with increasing frequency, as the signal-to-noise ratio becomes lower (see Methods, section 2.7).

In the process to detect local neural activity, we compared continuous multitaper and wavelet power estimation in two task-relevant regions from subject 1. In general, continuous multitaper estimation yielded smoother estimates, as expected. Fig. S1 (top and middle rows) compared spectrograms obtained with both techniques in a contact placed in V1/V2 and in a contact placed in the middle portion (anterior-posterior axis) of the MTG (mMTG). Continuous multitaper estimation revealed consistent high-gamma activity (64-256 Hz) from 300 to 600 ms after stimulus presentation in V1/V2, that was only barely detected by wavelet estimation (64-128 Hz, 400 to 500 ms). In the mMTG, the continuous multitaper spectrogram revealed more diffuse high-gamma activity (64-128 Hz) showing a special consistency from 500 to 800 ms after stimulus presentation. In contrast, those activations were not captured by wavelet estimation.

When averaging across trials, the high temporal resolution of wavelet estimation might cancel out high-frequency activations that are not homogeneously time-locked to the stimulus. To assess whether reduced sensitivity of wavelet estimation to high-frequency activity resulted from this effect, we also low-pass filtered the power estimates below 8 Hz before obtaining median spectrograms across trials (Fig. S1 bottom row). With this approach, high-gamma activations in V1/V2 revealed higher consistency and became comparable to those obtained with continuous multitaper estimation (64-256 Hz) from 400 to 500 ms. In contrast, high-gamma activations in the mMTG could not be recovered via wavelet esimation.

Hence, qualitative comparison of continuous multitaper and wavelet power estimation in two task-relevant regions from subject 1 (V1/V2 and mMTG) proved the multitaper method to be more sensitive to sustained high-frequency activations, while fast and highly variable fluctuations across trials tend to be attenuated. In particular, low-amplitude temporally diffuse high-gamma activity (64-128 Hz) in mMTG was found between 500 to 800 ms after stimulus presentation by the multitaper method, and, conversely, it could not be detected using the wavelet transform. Another caveat when combining temporally-resolved methods such as the wavelet transform with trial averaging is that small temporal misalignments in power activations across trials might result in these activations cancelling out in the average spectrogram. As a result, stimulus-locked events that occur within a certain temporal window but are not strictly locked to a time point, might remain invisible to exploration. This limitation can be partially overcome by low-pass filtering power estimates (i.e. temporally smoothing) before averaging across trials, as seen in Fig. S1 (bottom, left).

In conclusion, the choice of either method is highly dependent on the research question at hand. In our case, multitaper estimation served the purpose of finding robust task-relevant activations, while cancelling out spurious short-lasting fluctuations. However, in other situations (for instance, when exploring HFO activity, (see (Arnulfo et al., 2015) for an example), wavelet-derived methods might be preferable to capture short transient effects.

### S2 Coupling between local activity and global connectivity fluctuations: circular time-shifted surrogates

To assess the degree of correlation between each contact’s local activity *L_k_*(*t*) and the global connectivity fluctuations measured by the proposed global variable *G*(*t*), we used the following procedure. Let *L_k_* = [*L*_*k*,1_,…, *L_k,P_*] be the sample observations of variable *L_k_*(*t*) within the time period of interest for each *k*, where *P* is the total number of time points in the time series signal. Likewise, let *G* = [*G*_1_,…, *G_P_*] be the sample observations of variable *G*(*t*) within the time period of interest. Then, the coupling of both variables was estimated as the Spearman correlation between each time series: *ρ_k_* = *ρ*(*L_k_, G*), where *ρ*(·) stands for the Spearman correlation operator.

To infer the significance of each estimation, we used the circular shift methodology, which preserved the local and global variables’ autocorrelation properties (around 200 ms by definition) while destroying their temporal alignment. For each statistical test (i.e., for each contact *k*),> we generated 10,000 independent circular time-shifted surrogates by resampling the local variable using the following procedure. An independent integer *p* was randomly generated within the interval [400, *P* – 400]. Note that time shifts are selected to be larger than the autocorrelation length (*p* = 400 corresponds roughly to 200 ms). Then, the circular shift was performed by moving the first p values of *L_k_* to the end of the time series, creating the surrogate sample 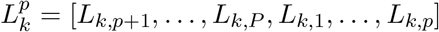. Then, we computed the Spearman correlation 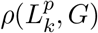 between the global time series and the local surrogate time series. By repeating this procedure for each circular timeshifted surrogate, we built a null distribution for the hypothesis that *L_k_* and *G* are uncorrelated. Only medium and large effect size correlations (*r* > 0.3) were kept. In addition, significance level was set to *α* = 0.05 and corrected for multiple comparisons using Bonferroni correction with the number of comparisons equal to the number of contacts.

**Figure S1:**
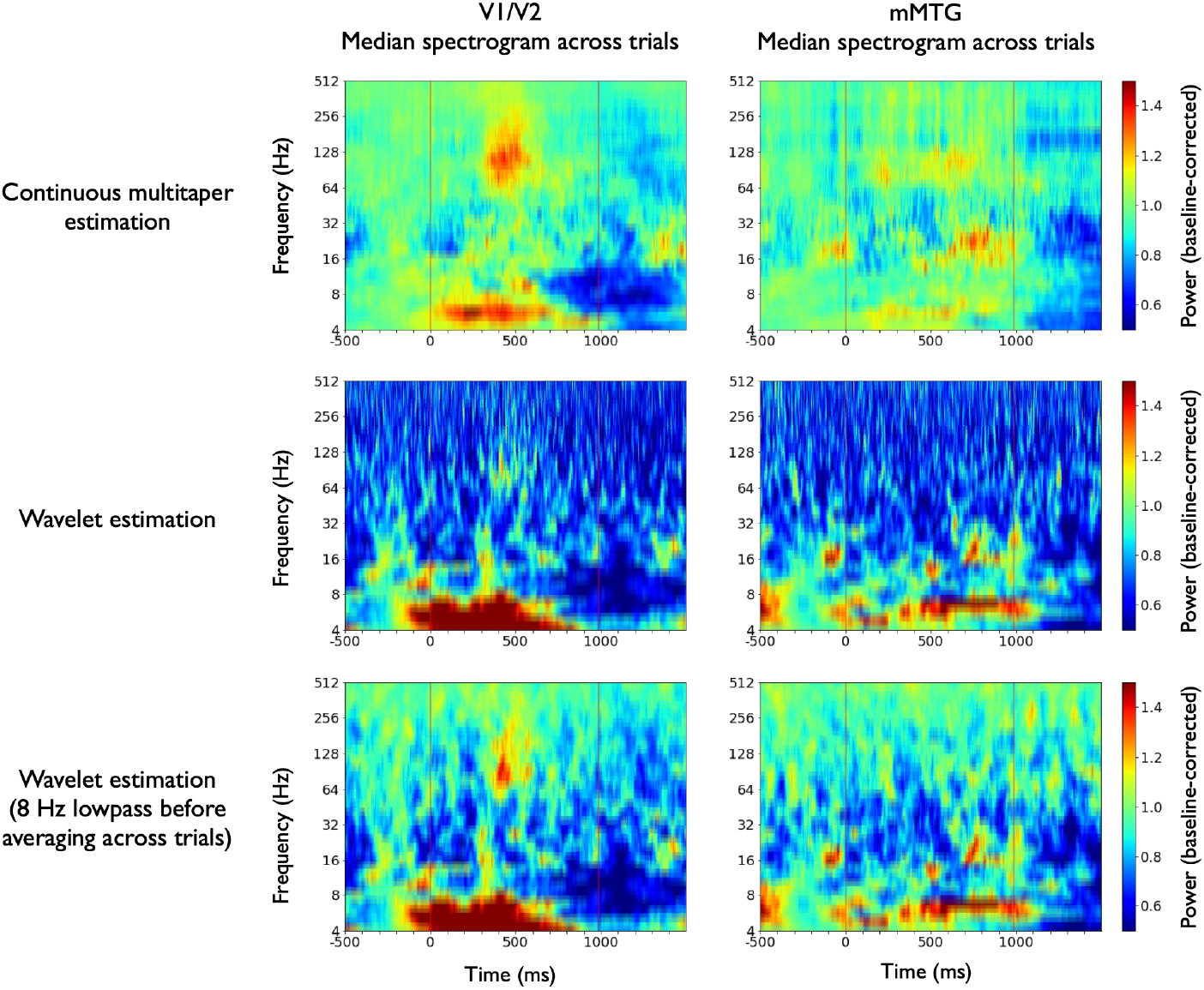
Continuous multitaper and wavelet spectral estimations for two exemplary task-relevant recording sites from subject 1. In general, multitaper spectrograms tend to be smoother and fast fluctuations are attenuated. In V1/V2 continuous multitaper estimation revealed consistent high-gamma activity (64-256 Hz) from 300 to 600 ms after stimulus presentation in V1/V2 (top left). These activations emerged, although less prominently with wavelet estimation (64-128 Hz, 400 to 500 ms, middle left). Low-pass filtering the wavelet power estimates below 8 Hz before obtaining the median spectrogram across trials turned to be more sensitive to these activations, due to potential time mis-alingments that were thus smoothed (bottom left). In the mMTG (middle portion of the MTG) the continuous multitaper spectrogram revealed diffuse high-gamma activity (64-128 Hz) with a special consistency from 500 to 800 ms after stimulus presentation. These activations were not captured by wavelet estimation. They also remained undetected after low-pass filtering the initial wavelet power estimates below 8 Hz.

**Figure S2:**
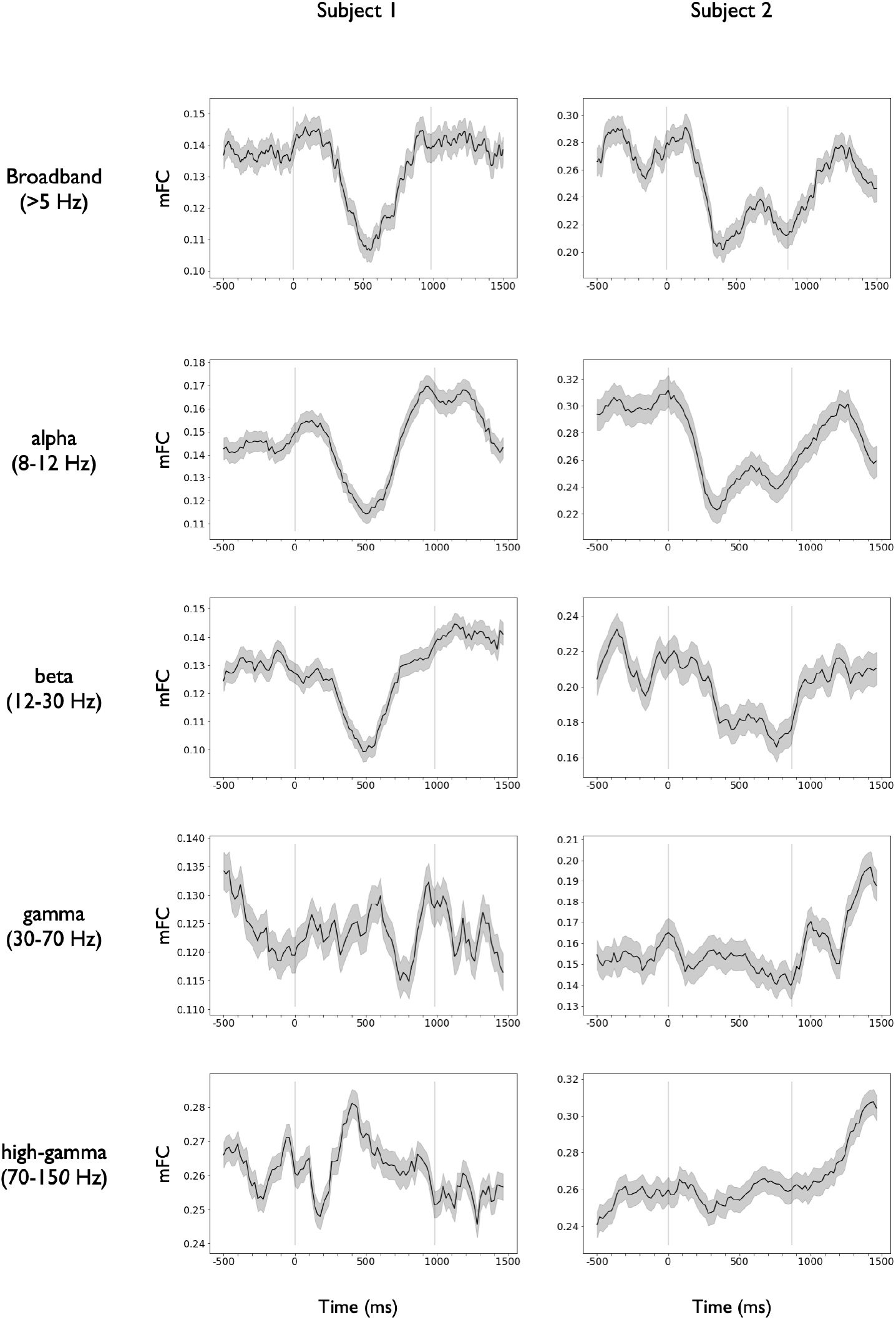
Decomposition of the mFC time course in its frequency components. The panel shows the time course of the mean functional connectivity (mFC) computed in the following bands: broadband (> 5 Hz), alpha (8-12 HZ), beta (12-30 Hz), gamma (30-70 Hz) and high-gamma (70-150 Hz). See Methods, section 2.8, for details. Vertical dark lines indicate the stimulus onset and offset times, respectively. A significant decrease in the mFC associated to stimulus presentation was found in the two subjects with respect to their pre-stimulus mean mFC value. This decrease was localized in the alpha and beta bands. Mean ± SEM across all GM contact pairs. Correlation values were Fisher’s z transformed before taking averages across contacts pairs.

**Figure S3:**
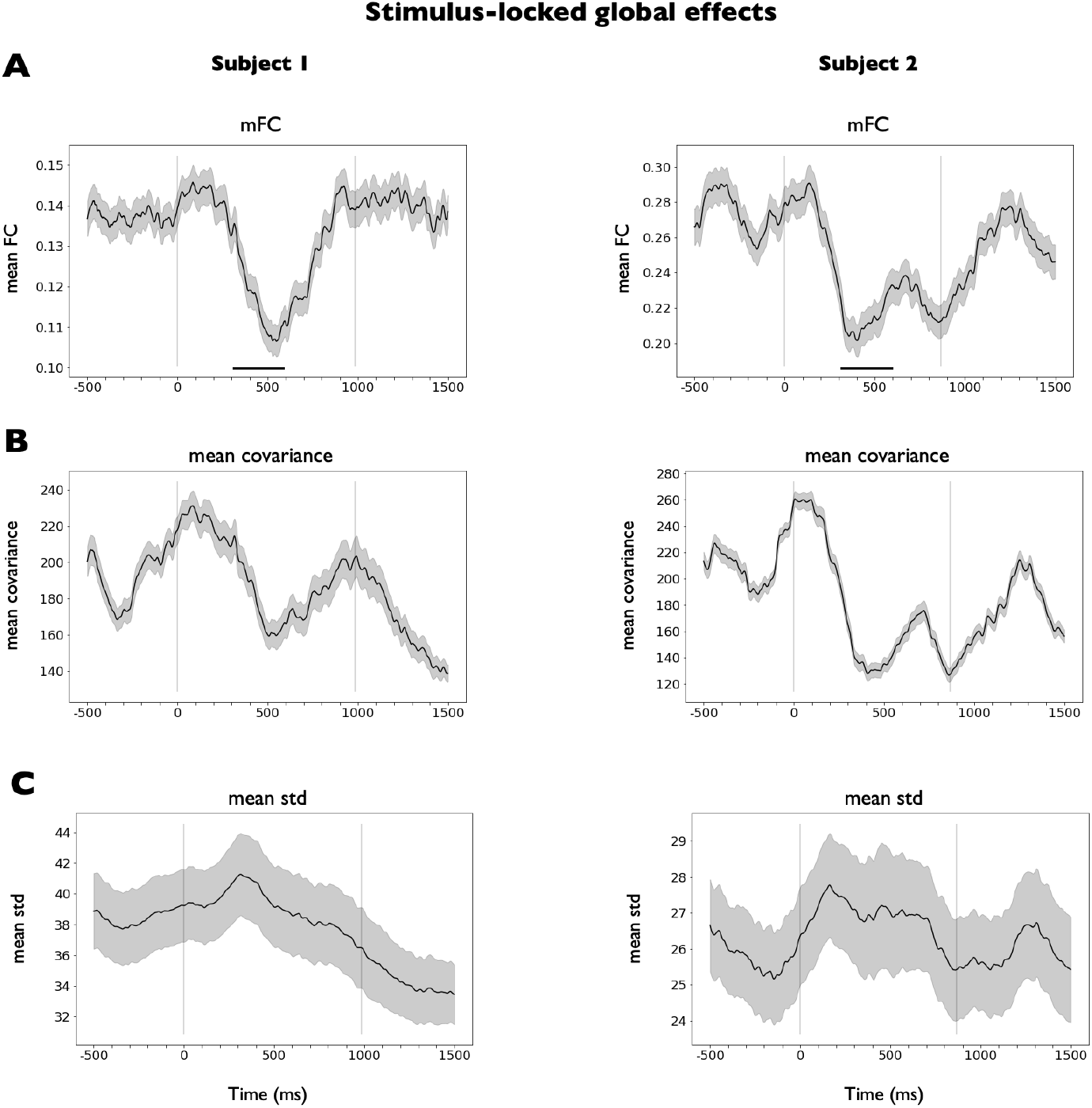
Dissection of the stimulus-locked global effects measured with mFC into its two components. **(A)** Time course of the mean functional connectivity (mFC). Mean ± SEM across all GM contact pairs (*N*_pairs_ = 1031 in subject 1, *N*_pairs_ = 307 in subject 2). Correlation values were Fisher’s z transformed before taking averages across contacts pairs. See Methods, section 2.8, for details. Vertical dark lines indicate the stimulus onset and offset times, respectively. A significant decrease in the mFC associated to stimulus presentation was found in the two subjects with respect to their pre-stimulus mean mFC value (300-600 ms, marked with black bars, Ranksum test applied across contact pairs on the average FC in the baseline period and the 300-600 ms window, *P* < 10^-6^, Cohen’s D effect size *D* > 5 in both subjects). **(B)** Time course of the average covariance across all pairs of selected GM contacts. Mean ± SEM across GM contact pairs (*N*_pairs_ = 1031 in subject 1, *N*_pairs_ = 307 in subject 2). **(C)** Time course of the average standard deviation across all GM contacts. Mean ± SEM across GM contacts (*N* = 47 in subject 1, *N* = 26 in subject 2).

**Figure S4:**
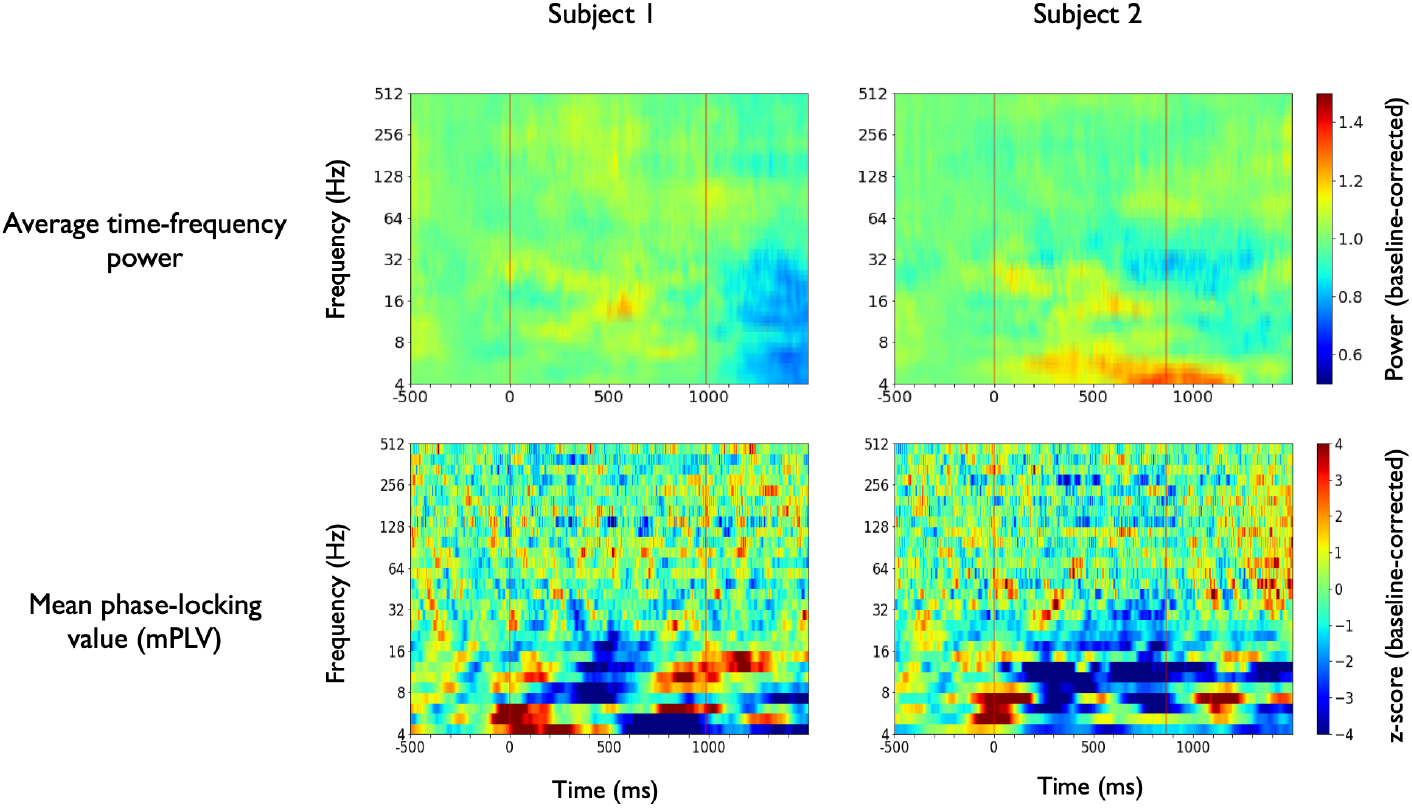
Comparison of average time-frequency power and mean phase-locking value (mPLV). **(top)** Average time-frequency power obtained by taking the mean spectrogram across all contacts. The figure has been baseline corrected (division by the mean power in the baseline period at each frequency). Vertical dark lines indicate the stimulus onset and offset times, respectively. The same scale is used as in S1. Only mild increases in global power are observed at around 500 ms after stimulus presentation in both patients. **(bottom)** Time-resolved mean phase locking value (mPLV) aligned to stimulus presentation. See Method, section 2.8, for details. Plots have been z-scored with respect to the pre-stimulus period (from −500 to −100 ms) at each frequency scale for visualization purposes. Vertical red lines indicate stimulus onset and offset times, respectively. A significant phase-decoupling was consistently observed across trials in the frequency range 6-16 Hz around 300 ms after stimulus presentation in the two subjects (z-score < 3). This inter-areal decoupling cannot be explained by a decrease in the signal-to-noise ratio or a simple decrease in power in 6-16 Hz, as there is no significant change in the global power.

**Figure S5:**
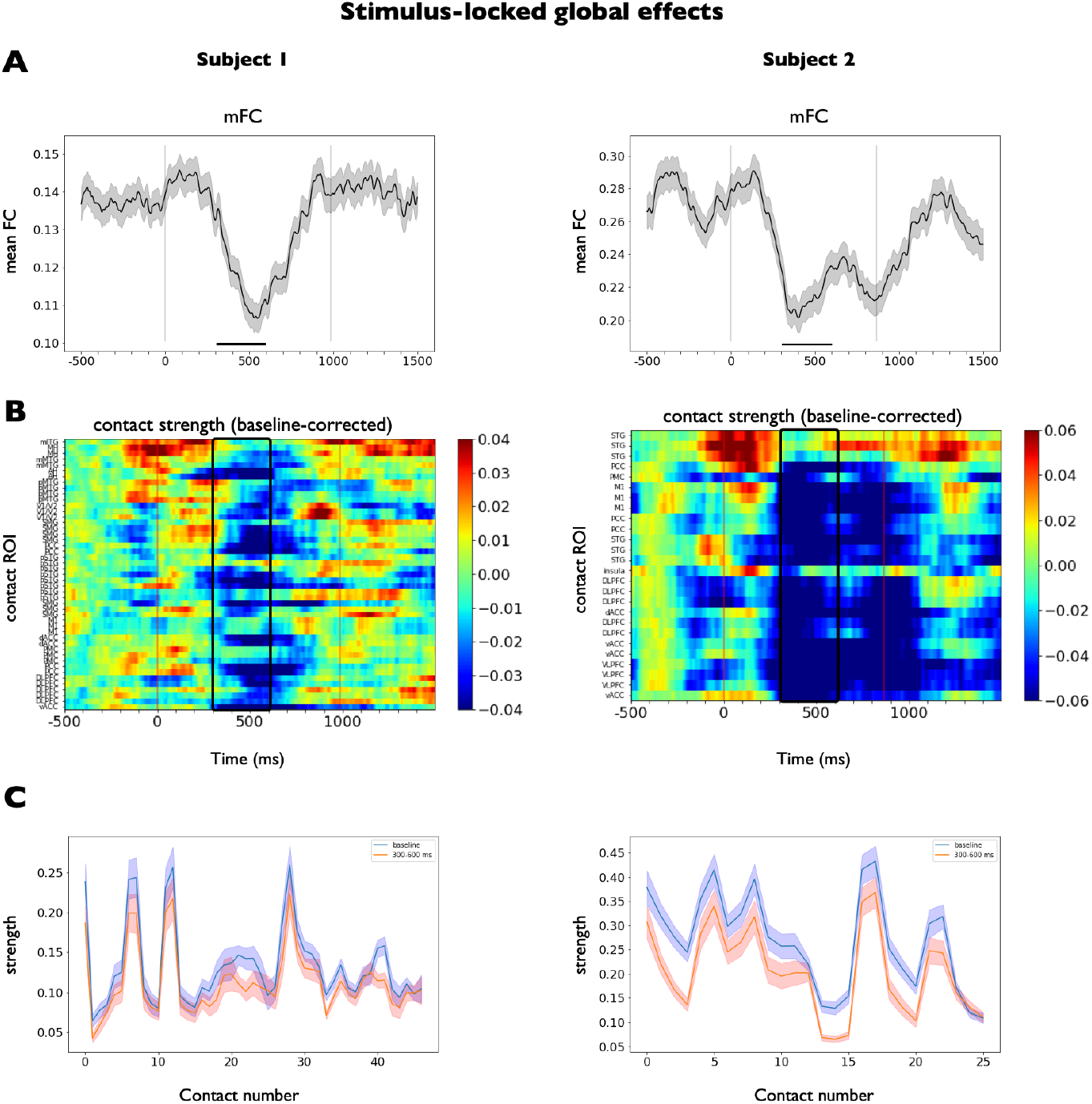
Connectivity fluctuations and contact strength across the task measured with FC. **(A)** Time course of the mean functional connectivity (mFC). Mean ± SEM across all GM contact pairs (*N*_pairs_ = 1031 in subject 1, *N*_pairs_ = 307 in subject 2). Correlation values were Fisher’s z transformed before taking averages across contacts pairs. See Methods, section 2.8, for details. Vertical dark lines indicate the stimulus onset and offset times, respectively. A significant decrease in the mFC associated to stimulus presentation was found in the two subjects with respect to their pre-stimulus mean mFC value (300-600 ms, marked with black bars, Ranksum test applied across contact pairs on the average FC in the baseline period and the 300-600 ms window, *P* < 10^-6^, Cohen’s D effect size *D* > 5 in both subjects). **(B)** Contact connectivity strength (average of the seed-based FC against its matching GM contact pairs). The period 300-600 ms is marked in red. For visualization purposes, strength values have been baseline-corrected by subtracting the average strength in the period from −500 to −200 ms before stimulus presentation (all windows fully contained in the baseline period). **(C)** Average contact strength in the pre-stimulus baseline period (from −500 to −200 ms, blue) and the marked stimulus epoch (300-600 ms). The decay in FC is a generalized effect across the implantation scheme of both subjects. Mean ± SEM across all contacts.

**Figure S6:**
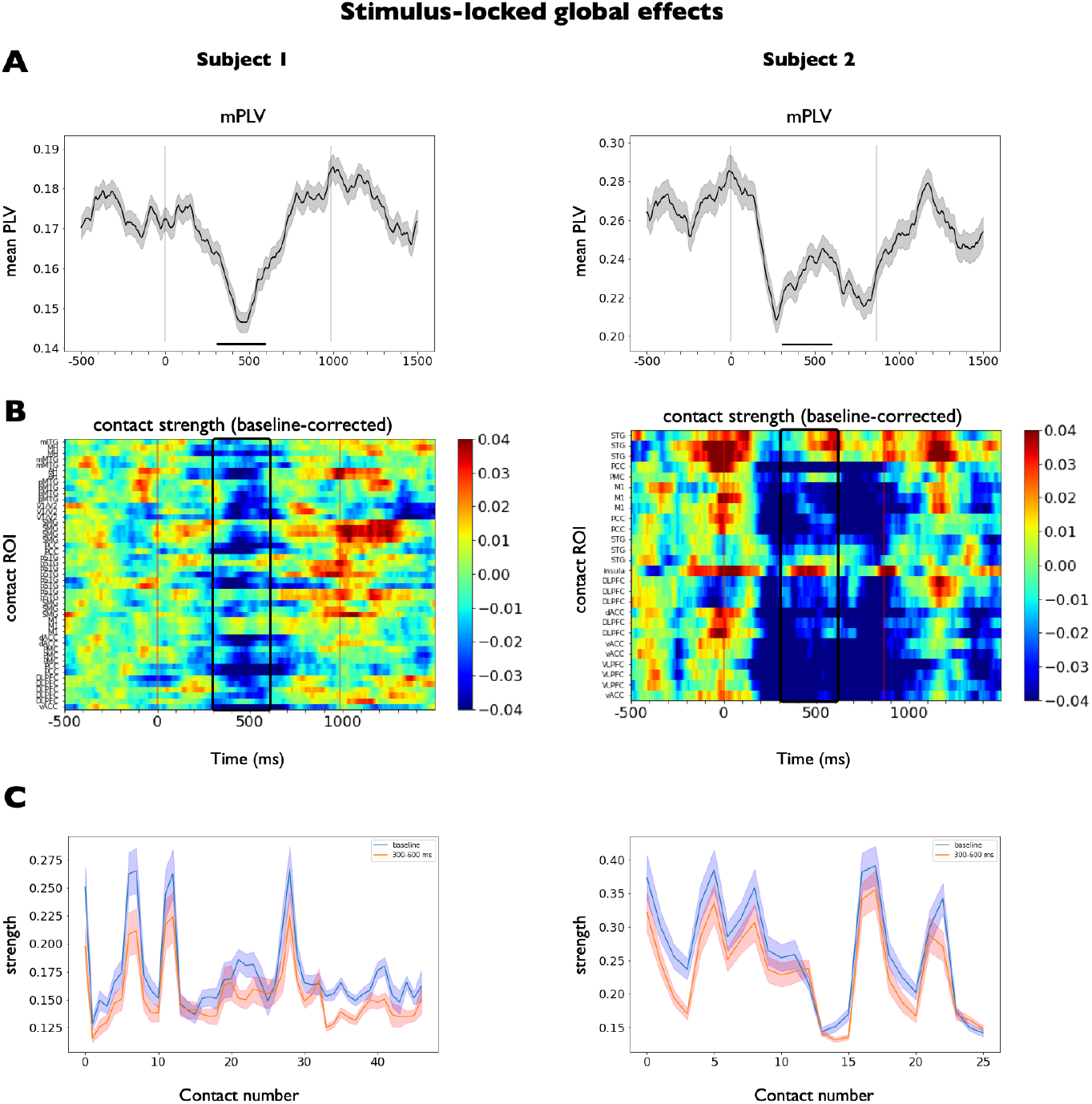
Connectivity fluctuations and contact strength across the task measured with PLV. **(A)** Time course of the mean phase-locking value (mPLV). Mean ± SEM across all GM contact pairs (*N*_pairs_ = 1031 in subject 1, *N*_pairs_ = 307 in subject 2). See Methods, section 2.8, for details. Vertical dark lines indicate the stimulus onset and offset times, respectively. A significant decrease in the mPLV associated to stimulus presentation was found in the two subjects with respect to their pre-stimulus mean mPLV value (300-600 ms, marked with black bars, Ranksum test applied across contact pairs on the average PLV in the baseline period and the 300-600 ms window, *P* < 10^-2^, Cohen’s D effect size *D* > 2.5 in both subjects). **(B)** Contact connectivity strength (average of the seed-based PLV against its matching GM contact pairs). The period 300-600 ms is marked in red. For visualization purposes, strength values have been baseline-corrected by subtracting the average strength in the period from −500 to −100 ms before stimulus presentation (all windows fully contained in the baseline period). **(C)** Average contact strength in the pre-stimulus baseline period (from −500 to −100 ms, blue) and the marked stimulus epoch (300-600 ms). The decay in PLV is a generalized effect across the implantation scheme of both subjects. Mean ± SEM across all contacts.

**Figure S7:**
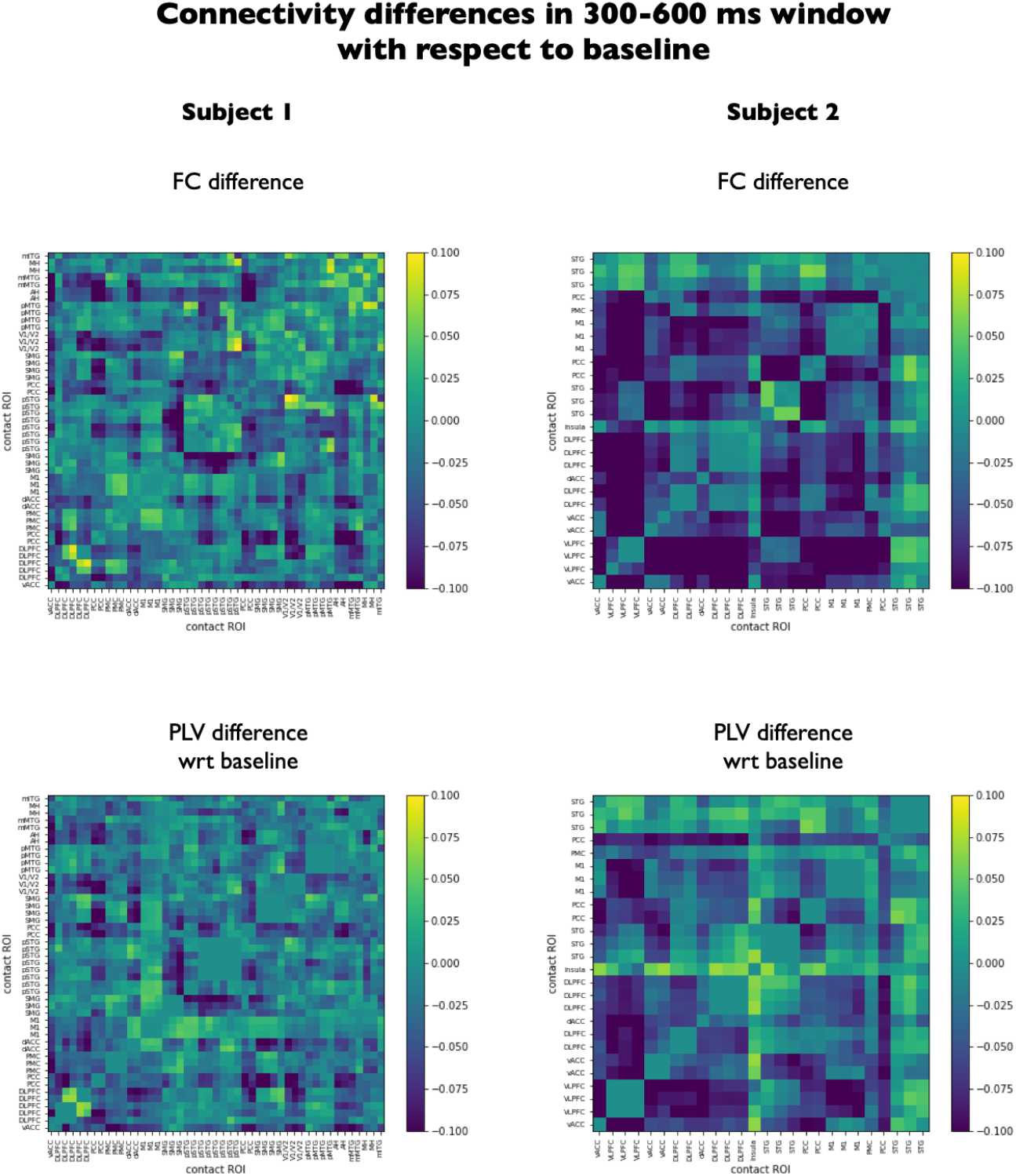
Connectivity differences in a 300-600 ms time-window with respect to baseline both with FC (top) and PLV (bottom). Average matrices were obtained in each period of interest. For each connectivity measure (FC and PLV) and subject (1 and 2), an average connectivity matrix was obtained both in the baseline period (from −500 to −100 ms) and in a post-stimulus period (300-600 ms). Here, we show the difference between the poststimulus matrix and the baseline matrix. The decay in FC and PLV is a generalized effect across most contact pairs for both subjects.

**Figure S8:**
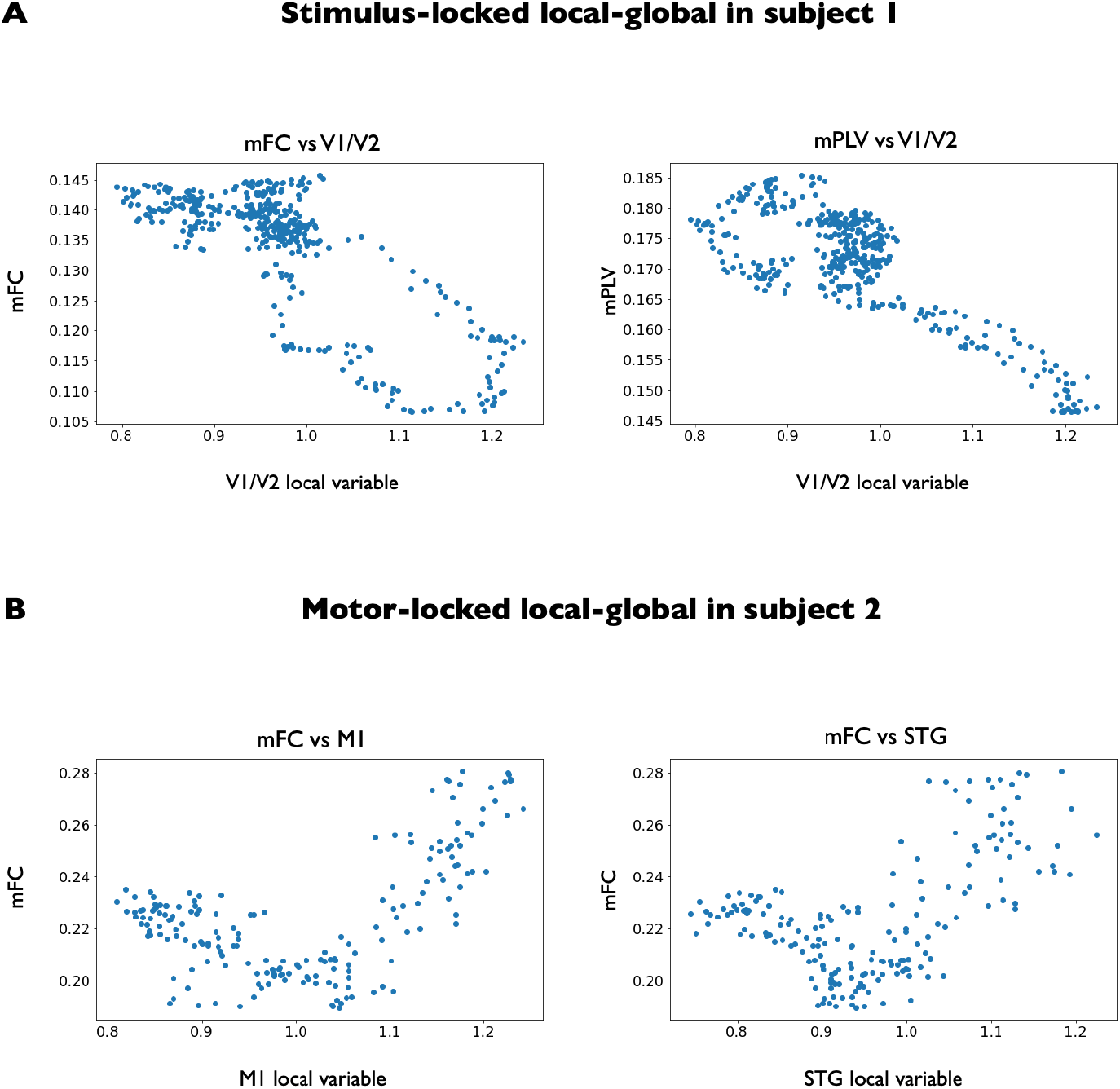
Exemplary scatter plots showing the association between local and global variables that were found to be significant. **(A)** Scatter plots in the stimulus-locked setting for subject 1 showing the association between mFC and a contact in V1/V2 (left), and between mPLV and a contact in V1/V2 (right). **(B)** Scatter plots in the motor-locked setting for subject 2 showing the association between mFC and a contact in M1 (left), and between mFC and a contact in STG (right). All variables have been downsampled by a factor of 10, to avoid autocorrelation effects.

**Table S1:** Demographic data. F: female

| Subject | Sex | Age<br>(years) | Epilepsy<br>onset<br>(years) | Suspected<br>epileptic<br>focus | Hand<br>laterality | Implantation | Number of<br>electrodes<br>(right/left) | Number of<br>contacts<br>(right/left) | Number of<br>contacts<br>included<br>in analysis |
| --- | --- | --- | --- | --- | --- | --- | --- | --- | --- |
| 3 | F | 23 | 7 | Bilateral<br>temporo-occipital<br>(heterotopia) | Right | Bilateral | 3/10 | 26/121 | 53 |

**Table S2:**
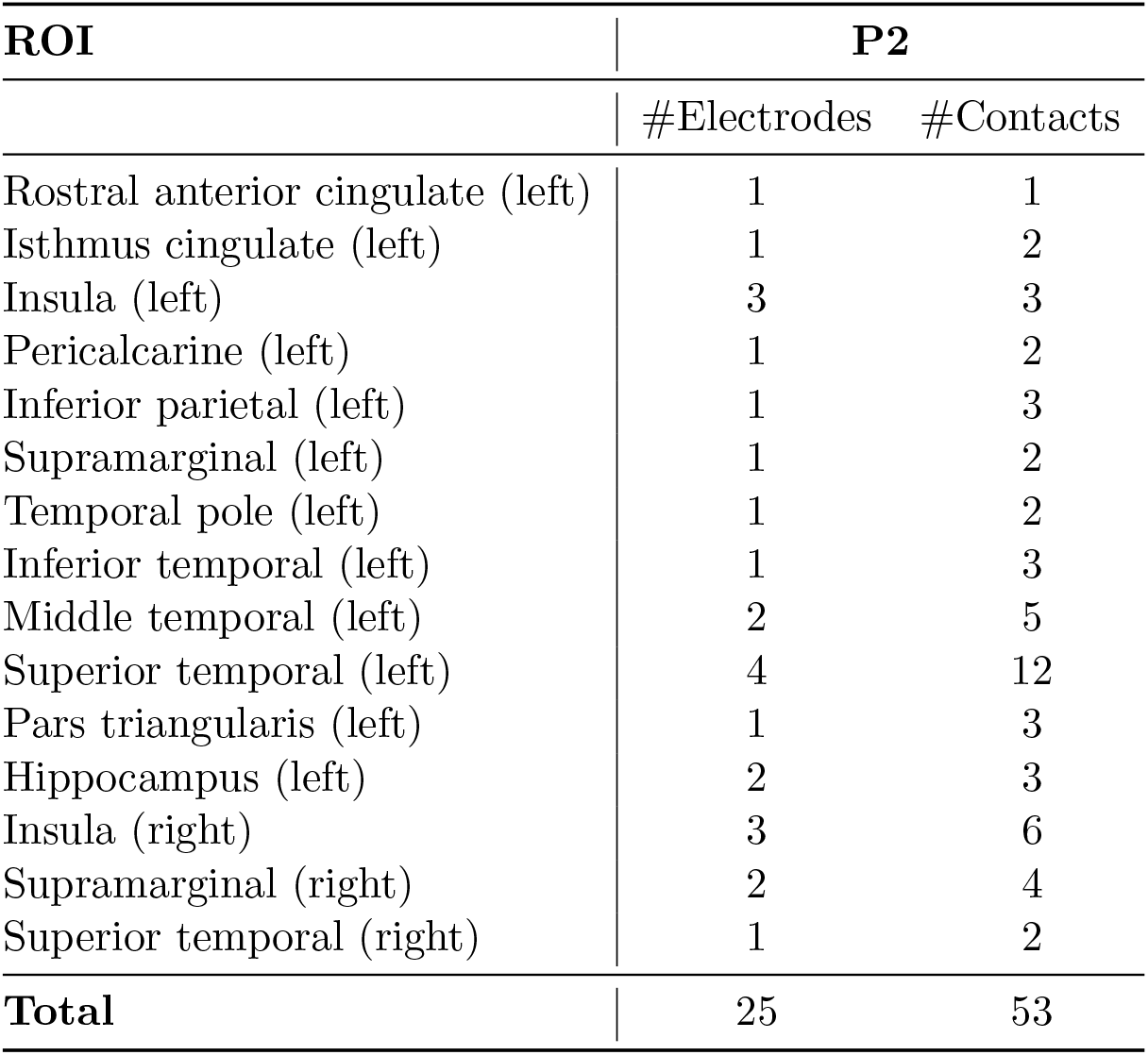
Implantation scheme. Regions of interest monitored in patient 3 are expressed in terms of the Desikan-Killiany atlas with an extra ROI for the hippocampus.

| ROI | P2 |  |
| --- | --- | --- |
|  | #Electrodes | #Contacts |
| Rostral anterior cingulate (left) | 1 | 1 |
| Isthmus cingulate (left) | 1 | 2 |
| Insula (left) | 3 | 3 |
| Pericalcarine (left) | 1 | 2 |
| Inferior parietal (left) | 1 | 3 |
| Supramarginal (left) | 1 | 2 |
| Temporal pole (left) | 1 | 2 |
| Inferior temporal (left) | 1 | 3 |
| Middle temporal (left) | 2 | 5 |
| Superior temporal (left) | 4 | 12 |
| Pars triangularis (left) | 1 | 3 |
| Hippocampus (left) | 2 | 3 |
| Insula (right) | 3 | 6 |
| Supramarginal (right) | 2 | 4 |
| Superior temporal (right) | 1 | 2 |
| **Total** | **25** | **53** |

**Figure S9:**
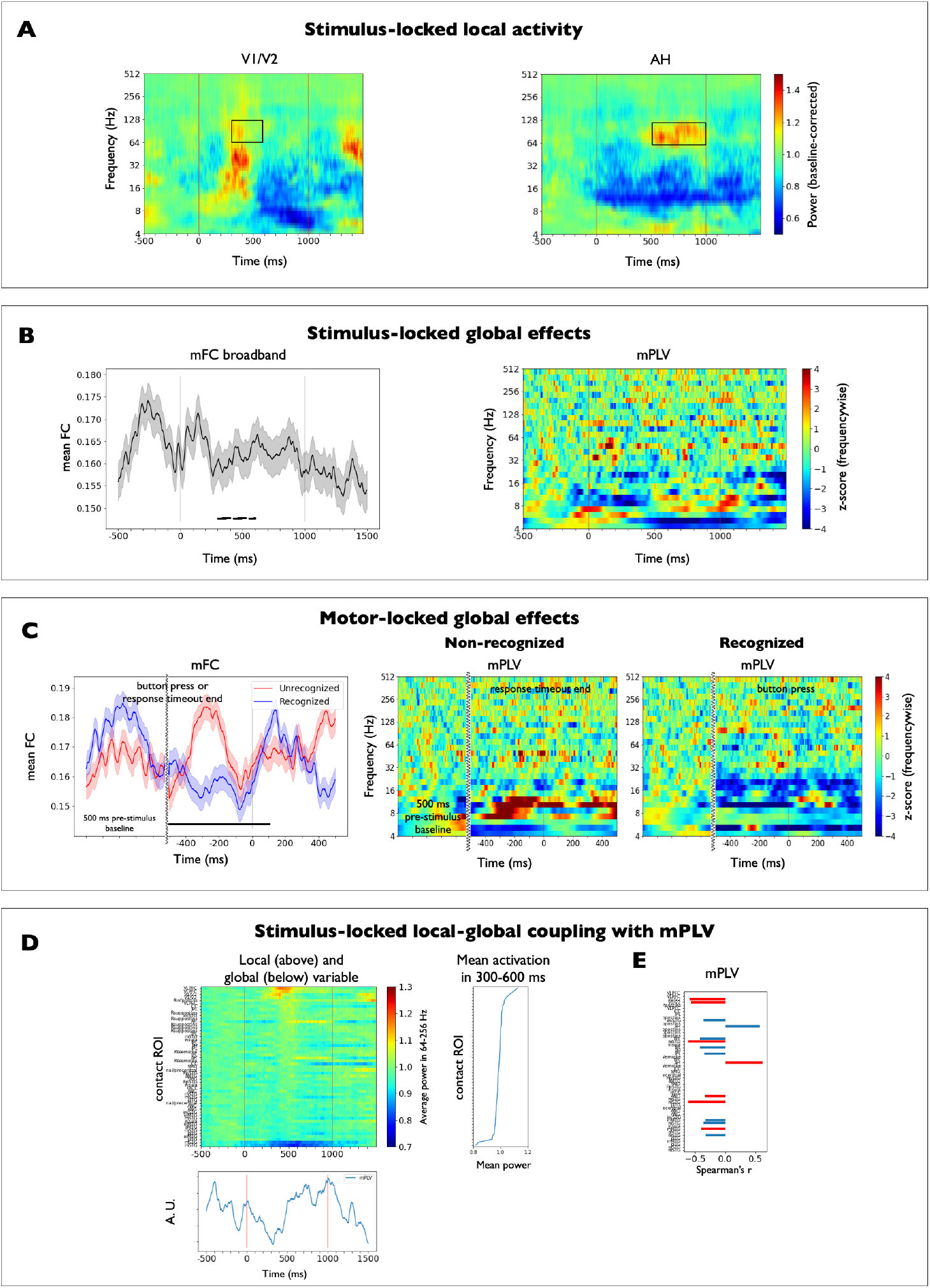
Cross-validation of local-global analysis on subject 3. **(A)** Stimulus-locked local activity. Median across all trials of the baseline-corrected spectrograms aligned to stimulus presentation (0 ms) in two exemplary recording sites of subject 3. Vertical lines indicate stimulus onset and offset times, respectively. Time-frequency windows of interest (TFOIs) used for statistical comparisons are marked with black rectangles. (Left) High-gamma activation of the visual cortex (V1/V2, 64 – 128 Hz, 300 – 600 ms, *P* < 10^-3^). (Right) High-gamma activation of the anterior hippocampus (AH, 64 – 128 Hz, 500 – 1000 ms, *P* < 10^-4^). **(B)** (Left) Stimulus-locked global effects measured with the time-resolved mean functional connectivity (mFC). Mean ± SEM across all GM contact pairs (N_pairs_ = 1341). Correlation values were Fisher’s z transformed before taking averages across contacts pairs. A noisy decreasing trend in the mFC associated to stimulus presentation was observed with respect to the pre-stimulus value. This trend was non-significant (300-600 ms, marked with a black dashed bar, Ranksum test applied across contact pairs on the average FC in the baseline period and the 300-600 ms window). (Right) Stimulus-locked global effects measured with the time-resolved mean phase locking value (mPLV). Values have been z-scored with respect to the pre-stimulus period (from −500 to −100 ms) at each frequency scale. A global phase-decoupling was found 300 ms after stimulus presentation in the frequency range 6-16 Hz (z-score < 3), similar to the trend observed in the primary analysis. **(C)** (Left) Motor-locked global effects measured with the broadband mFC in recognized (*N* = 51, blue line) and non-recognized (N = 45, red line) trials. Mean ± SEM across all GM contact pairs for each set of trials aligned to button press / response timeout end (marked with a vertical dark line), respectively. Pre-stimulus baseline is also shown for comparison. Curvy lines mark a discontinuity in time. A significant mFC difference between conditions was found between 500 ms before and 100 ms after button press or response timeout end, respectively (marked with a black bar, Ranksum test applied across contact pairs on the average FC in the selected time window across conditions, *P* < 0.05, Cohen’s D effect size *D* > 2). Note also the increase in mFC after button press, similarly found in the primary analysis. (Right) Motor-locked global effects measured with the mPLV in recognized (N = 51) and non-recognized (N = 45) trials, aligned to button press or response timeout end, respectively. **(D)** The upper panel shows the local variable for each brain recording site aligned to motor report (median spectrogram power across recognized trials aligned to button press, averaged across the high-gamma range, 64-256 Hz). Contacts have been sorted by the mean high-gamma power in the 300–600 ms time window after stimulus presentation, which defines the task-related activation of each contact (shown next to each contact’s power evolution). The lower panel shows the global variable mPLV, which was found to be modulated by the task. **(E)** Spearman correlation coefficient between local and global variables. Correlations have been thresholded at medium-size effects (*r* > 0.3). Significance was tested using a surrogate distribution via circular shifts, with a criterion of *α* = 0.05 and corrected for multiple comparisons for the number of contacts. Significant correlations are indicated with red bars. Local increase in activity at V1/V2 and STG contacts appears to have a significant correlation of −0.6 with the decrease in mPLV. These observations are consistent with the primary analysis. In addition, the decrease in mPLV seems to have a significant positive correlation with activity in AH.

## Footnotes

1 Manel Vila-Vidal

2 Mariam Khawaja

